# Fortuitous Somatic Mutations during Antibody Evolution Endow Broad Neutralization against SARS-CoV-2 Omicron Variants

**DOI:** 10.1101/2022.12.12.520172

**Authors:** Jianbo Wu, Zhenguo Chen, Yidan Gao, Zegen Wang, Jiarong Wang, Bing-Yu Chiang, Yunjiao Zhou, Yuru Han, Wuqiang Zhan, Minxiang Xie, Weiyu Jiang, Xiang Zhang, Aihua Hao, Anqi Xia, Jiaying He, Song Xue, Christian T. Mayer, Fan Wu, Bin Wang, Lunan Zhang, Lei Sun, Qiao Wang

**Affiliations:** Key Laboratory of Medical Molecular Virology (MOE/NHC/CAMS), Shanghai Institute of Infectious Disease and Biosecurity, Shanghai Frontiers Science Center of Pathogenic Microbes and Infection, Shanghai Fifth People’s Hospital, Shanghai Key Laboratory of Medical Epigenetics, Institutes of Biomedical Sciences, School of Public Health, School of Basic Medical Sciences, Fudan University, Shanghai 200032, China; Advaccine Biopharmaceuticals Suzhou Co., Ltd, Suzhou, China; Fundamental Research Center, Shanghai Yangzhi Rehabilitation Hospital (Shanghai Sunshine Rehabilitation Center), School of Medicine, Tongji University, Shanghai 201619, China; Experimental Immunology Branch, Center for Cancer Research, National Cancer Institute, National Institutes of Health, Bethesda, MD 20892, USA

**Keywords:** broadly neutralizing antibody (bNAb), SARS-CoV-2, variant of concern, clonally-related antibody family, somatic hypermutation

## Abstract

Striking antibody evasion by emerging circulating SARS-CoV-2 variants drives the identification of broadly neutralizing antibodies (bNAbs). However, how a bNAb acquires increased neutralization breadth during antibody evolution is still elusive. Here, we identified a clonally-related antibody family from a convalescent individual. One of the members, XG005, exhibited potent and broad neutralizing activities against SARS-CoV-2 variants, while the other members showed significant reductions in neutralization breadth and potency, especially against the Omicron sublineages. Structural analysis visualizing the XG005-Omicron spike binding interface revealed how crucial somatic mutations endowed XG005 with greater neutralization potency and breadth. A single administration of XG005 with extended half-life, reduced antibody-dependent enhancement (ADE) effect, and increased antibody product quality, exhibited a high therapeutic efficacy in BA.2- and BA.5-challenged mice. Our results provided a natural example to show the importance of somatic hypermutation during antibody evolution for SARS-CoV-2 neutralization breadth and potency.

## INTRODUCTION

Variant strains of severe acute respiratory syndrome coronavirus 2 (SARS-CoV-2) continue to emerge and spread globally. So far, five variants of concern (VOCs) have been defined, including Alpha (B.1.1.7), Beta (B.1.351), Gamma (P.1), Delta (B.1.617.2), and the newly identified Omicron (B.1.1.529) variants (Karim and Karim, 2021; Mannar et al., 2022; Viana et al., 2022). These VOCs bear mutations in the viral spike protein (S protein), not only increasing the viral transmissibility or virulence, but also facilitating the immune escape (Altmann et al., 2021; Mlcochova et al., 2021; Planas et al., 2021; Wang et al., 2021a; Wang et al., 2021b). Many monoclonal antibodies (mAbs) identified from convalescent or vaccinated individuals showed diminished or abrogated neutralizing activity against distinct VOCs (Schmidt et al., 2021; Wang et al., 2022b). Especially, the newly emerged Omicron variant encodes 37 amino acid substitutions in the viral S protein, 15 of which are located in the receptor-binding domain (RBD), and causes significant humoral immune evasion, posing a remarkable challenge for the effectiveness of vaccines and mAb therapies (Cameroni et al., 2022; Cao et al., 2022a; Carreno et al., 2022; Cele et al., 2022; Iketani et al., 2022; Liu et al., 2022; Planas et al., 2022; Zhou et al., 2022).

These newly emerging SARS-CoV-2 variants with strong immune escape capacity motivate researchers to identify broadly neutralizing antibodies (bNAbs) that could be of potential clinical benefit. Combining two mAbs recognizing two distinct epitopes is a popular strategy to increase the neutralizing breadth and avoid viral evasion (Baum et al., 2020; Dong et al., 2021; Li et al., 2022). For example, Eli Lilly’s combination of two RBD-binding mAbs, bamlanivimab (LY-CoV555) and etesevimab, has been authorized for emergency use after exposure to the SARS-CoV-2 virus (Dougan et al., 2021). Tixagevimab (AZD8895) and cilgavimab (AZD1061) combination showed both prophylactic and therapeutic efficacy in a nonhuman primate model of SARS-CoV-2 infection (Loo et al., 2022). A bispecific antibody through connecting two single-domain antibodies, n3113v and n3130v, also displayed exceptional neutralizing breadth and potency via inhalation administration (Li et al., 2022).

Meanwhile, using just a single monoclonal bNAb with high neutralization potency and breadth could also be effective for clinical prevention or therapy. For example, LY-CoV1404 (also known as bebtelovimab) exhibits exceptional neutralizing activity against various SARS-CoV-2 variants, unaffected by most of these variant mutations (Iketani et al., 2022; Westendorf et al., 2022; Zhou et al., 2022). However, the number of super-antibodies with extreme broad-spectrum activity and ultra-potency is still very limited, and more importantly, its evolution process in vivo is still largely unknown.

Here, we screen mAbs isolated from a convalescent donor with elite serum neutralizing activity (Zhou et al., 2021), and identified XG005, a fully human IgG1 mAb targeting SARS-CoV-2 RBD, as an extremely potent neutralizing antibody, both in vitro and in vivo, against all currently known VOCs and the most recently emerged Omicron variants, BA.1, BA.2, BA.2.12.1, BA.3, and BA.4/5, which have severe immune escape capacity (Cao et al., 2022a; Cao et al., 2022b; Iketani et al., 2022; Liu et al., 2022). Structural analysis revealed that XG005 bound to an epitope that overlapped with VOC escape mutations, but delicately avoided immune escape and retained its binding affinity. Moreover, three clonally-related family members of XG005 isolated from the expanded B cell clone of the same donor showed reduced levels of neutralizing potency and breadth, suggesting that the resistance of XG005 evolved stochastically. Comparison of their sequences identified the somatic mutations at the amino acid residues crucial for antibody neutralizing potency and breadth. Considering that this convalescent individual donated the blood at a time when there were no emerging variants of SARS-CoV-2, we conclude that a highly potent and broad neutralizing antibody could evolve stochastically even in convalescent individuals whose sera barely neutralize SARS-CoV-2 Omicron variants.

## RESULTS

### Screening of antibodies isolated from a convalescent donor

We isolated monoclonal antibodies (mAbs), XG001-XG048, from a convalescent individual who donated blood in April 2020 when no SARS-CoV-2 variant had been reported (Zhou et al., 2021). Half of these antibodies (23/45, red name in Figure 1A) recognized the receptor-binding domain of SARS-CoV-2 spike protein (S protein); one fourth (11/45, blue name in Figure 1A) were N-terminal domain (NTD)-binding antibodies; and several (5/45, green name in Figure 1A) bound S2 stalk region (Zhou et al., 2021). To explore the cross-reactivity of these antibodies against different VOCs, we first performed an ELISA analysis against the S-protein of SARS-CoV-2 and its related VOCs (Figure 1A). Among 45 antibodies, 2, 8, 5, 7 and 23 antibodies exhibited at least 25% reduction of binding activity against S protein of B.1.1.7 (Alpha), B.1.351 (Beta), P.1 (Gamma), B.1.617.2 (Delta) and B.1.1.529 (Omicron) variants, respectively (Figure 1B). Some antibodies, such as XG027 and XG043, showed a substantial loss in antigen binding against most VOCs; for some others, such as RBD-binding antibody XG005 and NTD-binding antibody XG035, no loss of binding capacity was observed. Together, these results suggest that Omicron exhibited a higher level of resistance to the tested mAbs isolated from a convalescent individual, and that many mAbs maintain their binding capacity against VOCs.

**Figure 1.**
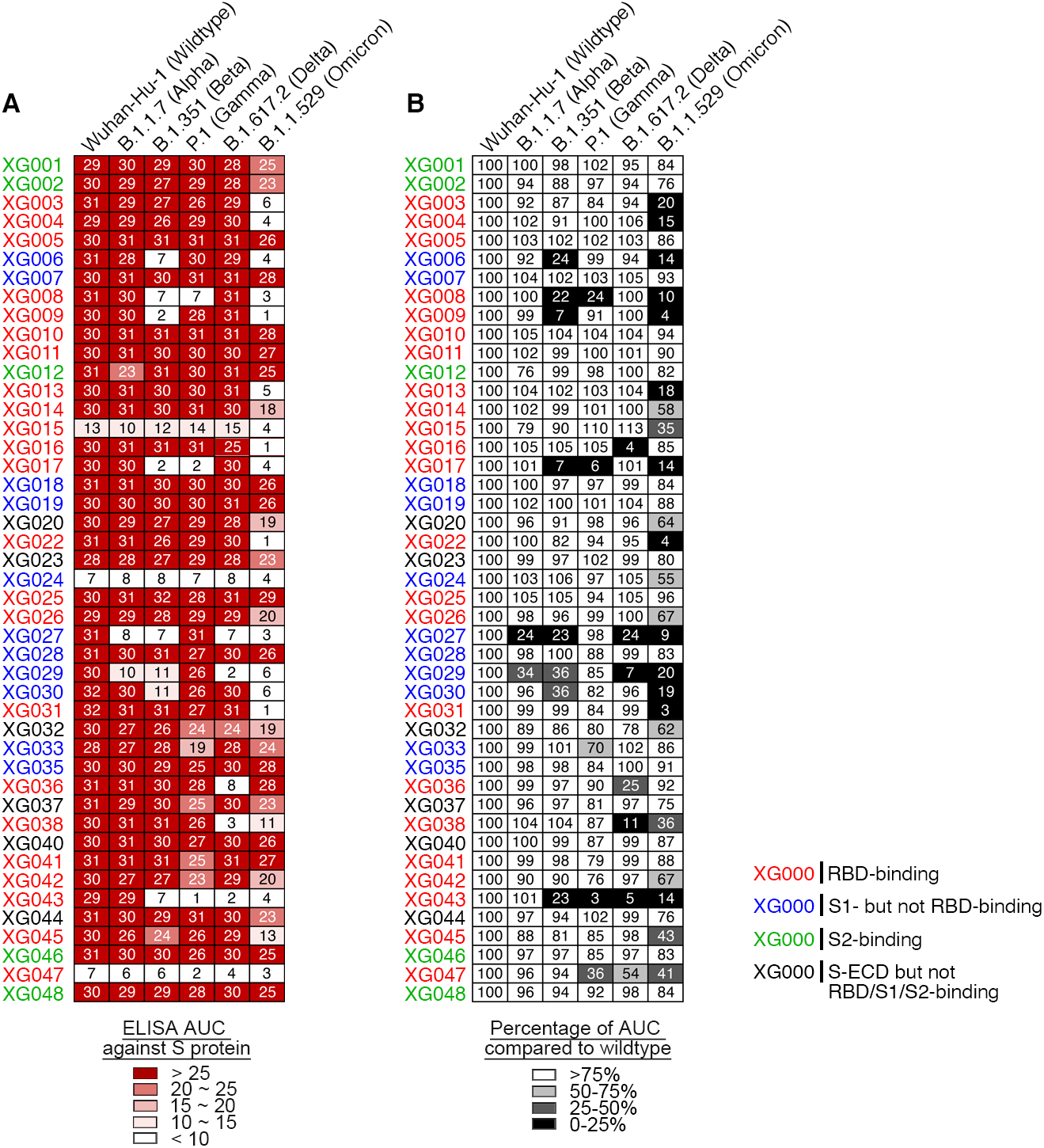
ELISA cross-reactivity of anti-S monoclonal antibodies. (A) Graphs show antibody ELISA reactivity against S proteins of wild-type SARS-CoV-2 and its five related VOCs. The six tested ELISA antigens include S proteins of Wuhan-Hu-1 (wild-type), B.1.1.7 (Alpha), B.1.351 (Beta), P.1 (Gamma), B.1.617.2 (Delta) and B.1.1.529 (Omicron). ELISA area under the curve (AUC) values were calculated for all 45 monoclonal antibodies (mAbs) isolated from a convalescent donor with a potent serum neutralizing activity (Zhou et al., 2021). Representative of two experiments. The names of monoclonals are color-coded: red, RBD-binding mAb; blue, NTD-binding mAb; green, S2-binding mAb; and black, S-ECD- but not RBD/S1/S2-binding mAb. (B) Percentage change in ELISA AUC relative to wild-type S protein. ELISA AUC results are presented as percentage of AUC normalized to the reactivity against Wuhan-Hu-1 (wild-type) S protein, and are illustrated by colors: black, 0%–25%; dark gray, 25%–50%; light gray, 50%–75%; and white, >75%.

### Neutralizing activity *in vitro* against VOCs

Antibody binding cannot predict viral neutralization. To assess the neutralization profile of these mAbs, we constructed various luciferase-expressing SARS-CoV-2 pseudoviruses, including SARS-CoV-2 Wuhan-Hu-1 (wild-type), B.1.1.7 (Alpha), B.1.351 (Beta), P.1 (Gamma), B.1.617.2 (Delta) and B.1.1.529 (Omicron) variants, and performed *in vitro* neutralization assays and calculated the IC_50_ values (Liu et al., 2021; Zhou et al., 2021) (Figure 2A and 2B). Twenty-three antibodies were neutralizers against wild-type SARS-CoV-2, and all of them, except XG005, partially or entirely, lost their neutralizing activity to at least one VOC (Figure 2A and 2B). Some monoclonal antibodies, such as XG001 and XG002, were not neutralizing at all, while potent neutralizers XG014 and XG016 showed significant antibody evasion by only the Omicron variant (Figure 2C). XG005 exhibited ultra-potent neutralizing activities against all VOCs (Figure 2C).

**Figure 2.**
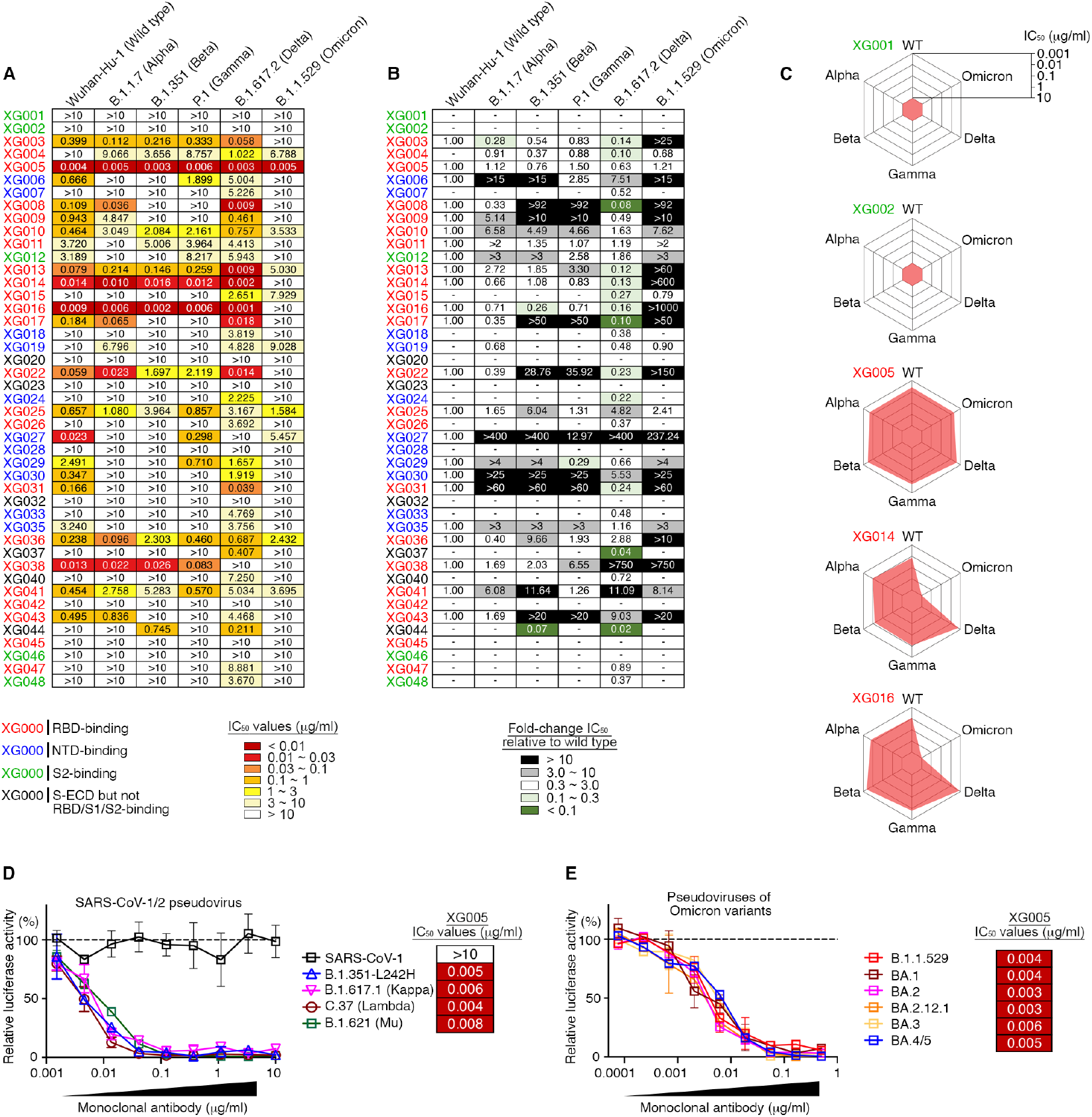
Cross-neutralization by monoclonal antibodies. (A) Pseudovirus neutralization assays by monoclonal antibodies. IC_50_ values for all 45 antibodies measured against Wuhan-Hu-1 (wild-type), B.1.1.7 (Alpha), B.1.351 (Beta), P.1 (Gamma), B.1.617.2 (Delta) and B.1.1.529 (Omicron) pseudoviruses. Antibodies with IC_50_ values above 10 μg/ml were shown as >10 μg/ml. Mean of two independent experiments. The names of monoclonals are color-coded: red, RBD-binding mAb; blue, NTD-binding mAb; green, S2-binding mAb; and black, S-ECD- but not RBD/S1/S2-binding mAb. (B) Fold change in IC_50_ values relative to Wuhan-Hu-1 (wild-type) SARS-CoV-2. Reduced neutralizing activities (increased IC_50_ values) are presented in black (>10-fold) or gray (3-10-fold), while enhanced neutralization (decreased IC_50_ values) in dark green (<10%) and light green (10-30%). (C) Spider charts for IC_50_ values of representative monoclonal antibodies. (D and E) Neutralization potency of XG005. Luciferase-based pseudoviruses of SARS-CoV-1, four SARS-CoV-2 variants (D) and six Omicron variants (E) were used for cell infection, and the luciferase signal after infection was determined as a surrogate of infection and normalized to the no antibody control (dashed line). In vitro neutralization assays for each antibody were performed at least two times, presented as mean ± SEM. IC_50_ values, mean of two independent experiments.

Consistent with other reports that the Omicron variant escapes antibody neutralization strikingly (Iketani et al., 2022; Liu et al., 2022), nearly 90% of the our neutralizing antibodies (20/23) had impaired Omicron neutralization with a more than 3-fold increase in the antibody IC_50_ values (Figure 2B). Among the 10 Omicron-neutralizing antibodies, 9 had IC_50_ values ranging from 1-10 μg/ml, and only one, XG005, exhibited an impressive neutralizing potency, with an IC_50_ value of 0.005 μg/ml (Figure 2A). Taken together, these results suggest that all tested mAbs isolated from this donor, except XG005, significantly lost their neutralizing activities against VOCs, especially against Omicron variants.

### Broad neutralizing activity of XG005

The outstanding neutralizing activity of XG005 led us to further assess the neutralization profile of XG005. We constructed several more types of pseudoviruses, including SARS-CoV-1, SARS-CoV-2 variants [B.1.351-L242H, B.1.617.1 (Kappa), C.37 (Lambda), B.1.621 (Mu)], and SARS-CoV-2 Omicron variants [BA.1, BA.2, BA.2.12.1, BA.3, BA.4/5], and performed pseudovirus neutralization assays. XG005 remained potent in neutralizing all these variants, including Omicron sublineages, with IC_50_ values of 0.008 μg/ml or lower, but had no neutralization activity against SARS-CoV-1 (Figure 2D and 2E). Together, the potent and broad neutralizing activity of XG005 indicates that there is still a highly conserved RBD epitope for antibody binding which is not affected by any escape mutations in SARS-CoV-2 variants.

### Structural and functional basis of XG005 neutralization and retained potency

To understand the structural basis for the neutralizing activity of XG005, we determined the cryo-EM structure of the SARS-CoV-2 wild-type S trimer complexed with XG005 Fab, revealing a conformation of two “up” and one “down” RBD with three Fabs (UDU with three Fabs, PDB ID 7V26, 3.8 Å resolution)(Liu et al., 2021). To further understand its broad neutralizing activity, we determined the cryo-EM structure of the SARS-CoV-2 Omicron S trimer complexed with XG005 (OS-XG005) (Table S1). Other than the UDU conformation with three Fabs (PDB ID 7YCZ, 3.24 Å), the OS-XG005 exhibited another two states, one “up” and two “down” RBDs with two Fabs (UDD with two Fabs, 3.62 Å), and one “up” and two “down” RBDs with three Fabs (UDD with three Fabs, PDB ID 7YCZ, 3.74 Å) (Figure 3A). Among these conformations, the “up” RBDs opened almost in the same orientation, while the orientations of “down” RBDs were different, which might result from the conformations of the other two RBDs (Figure 3B).

**Figure 3.**
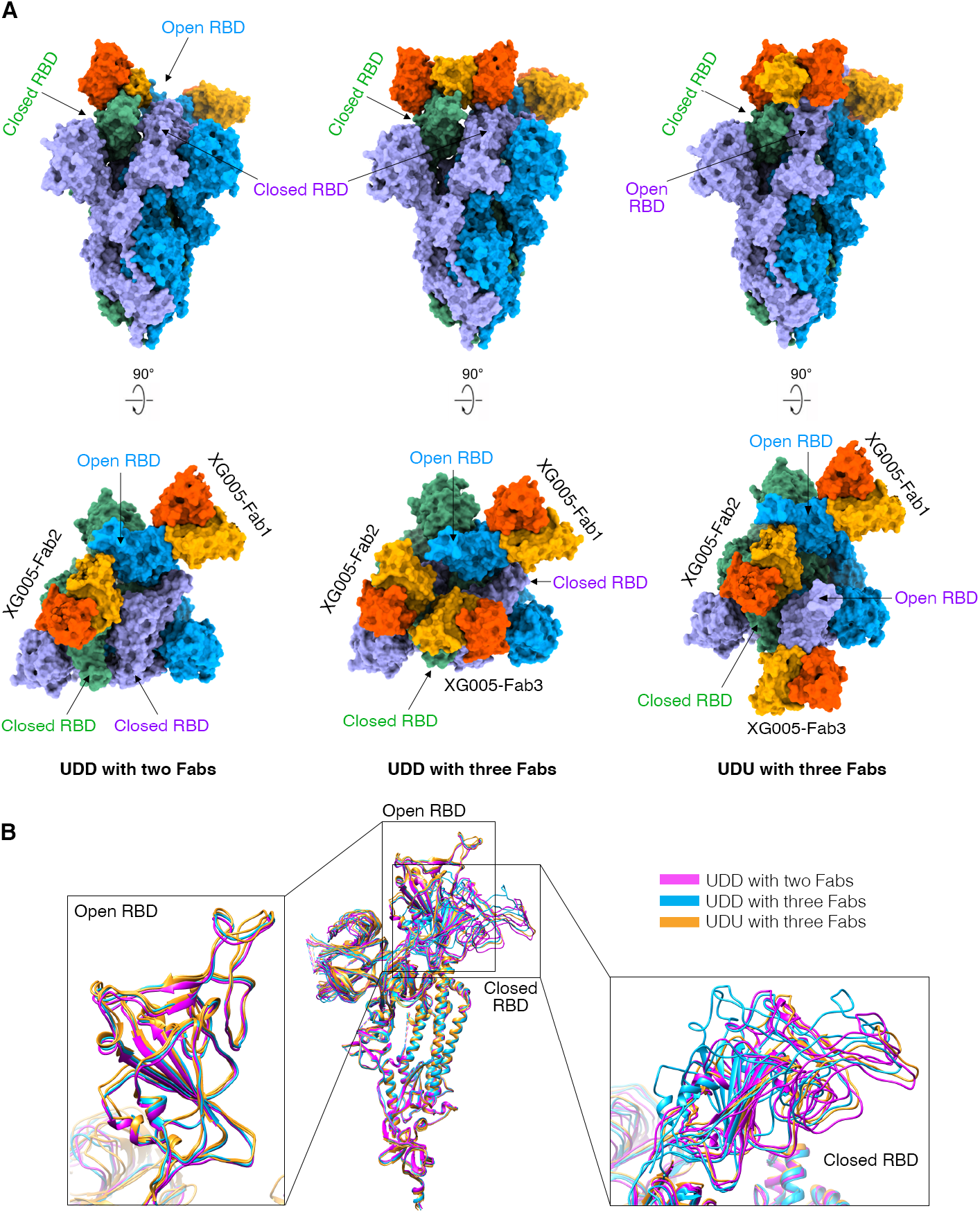
Cryo-EM structure of XG005 complexed with Omicron S trimer. (A) XG005 binds to Omicron S trimer in three states: one “up” RBD and two “down” with two Fabs (UDD with two Fabs), one “up” RBD and two “down” with three Fabs (UDD with three Fabs), and two “up” RBDs and one “down” RBD with three Fabs (UDU with three Fabs). Two perpendicular views of Omicron S-XG005 depict the surface. The XG005 VH/CH and VL/CL domains are colored in orange and yellow, respectively. Three S protomers of Omicron S trimer are colored in blue, green and purple, respectively. (B) Comparison of all S monomers of the three states in Ribbon, showing that all “up” RBDs are at the similar orientation while the down RBDs adopt different orientations. The monomers of three states are colored in magenta, blue and yellow, respectively.

Comparison of the interface regions of wild-type RBD-XG005 (Liu et al., 2021) and Omicron RBD-XG005 showed that the XG005 interacted with wild-type and Omicron RBD in a very similar way. The tight contacts between XG005 and Omicron RBD mainly resulted from extensive hydrophilic interactions. Three mutation residues (N440K, G446S and N501Y) of the Omicron S were located in the XG005-recognizing epitope (Figure 4A and 4B). Specifically, although N501Y mutation led to the loss of two hydrogen bonds between N501 of wild-type RBD and N33 of XG005 CDRL1 (Figure 4C and 4D), G446S mutation introduced two hydrogen bonds between Omicron-S S446 and T96 of XG005 CDRL3 (Figure 4C and 4D). Moreover, N440K mutation destroyed the hydrogen bonds between residues N440/L441 of wild-type RBD and Y34/G33 of XG005 CDRL1/CDRH2, but rescued one hydrogen bond between K440 of Omicron RBD and A103 of XG005 CDRH3 (Figure 4C and 4D). In addition, one hydrogen bond formed between N450 of Omicron RBD and D58 of XG005 CDRH2 as a compensation (Figure 4C and 4D). Therefore, the three Omicron mutations (N440K, G446S and N501Y) did not disrupt the RBD-XG005 interaction (Figure 4E).

**Figure 4.**
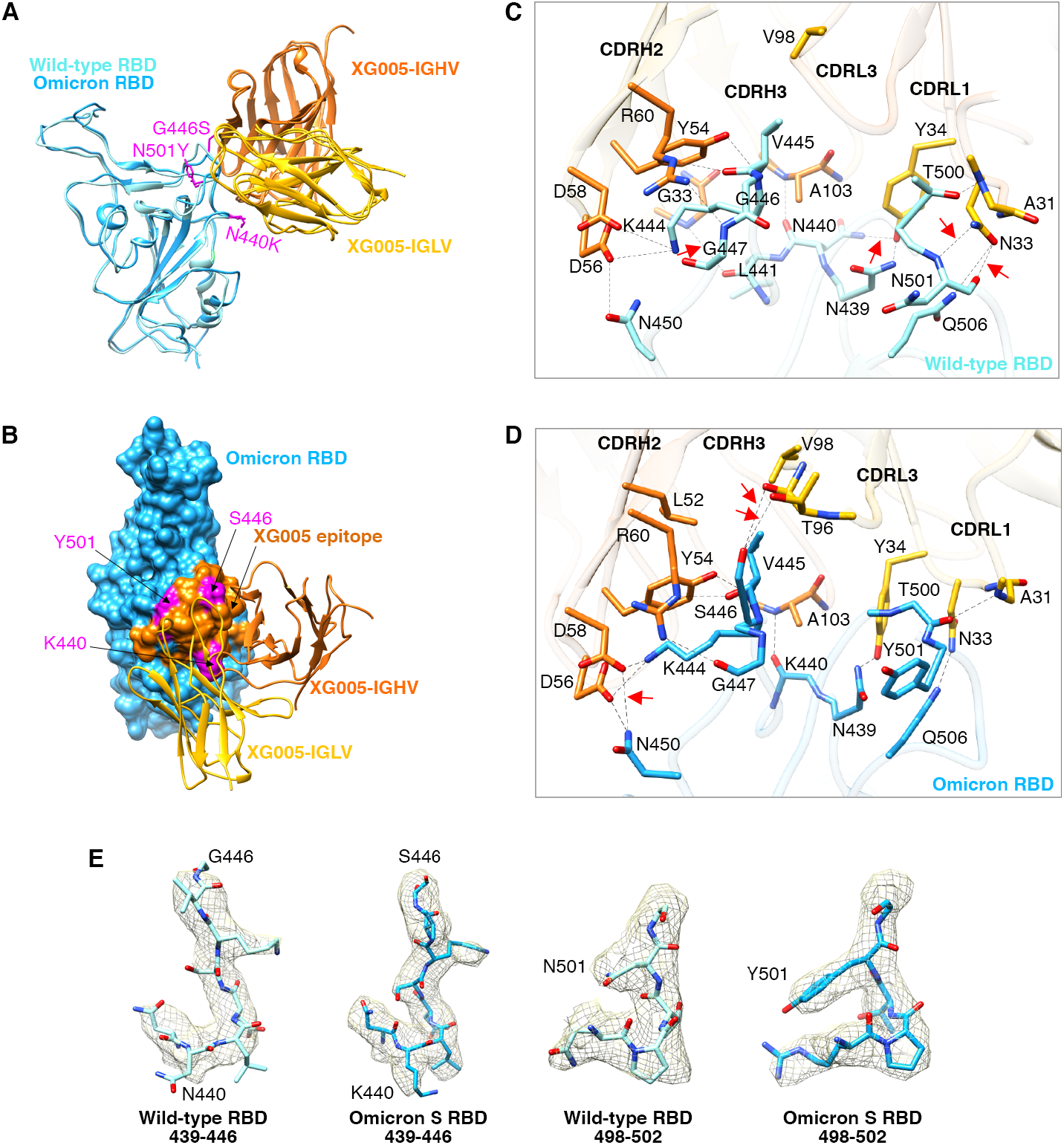
Comparison the interface between SARS-CoV-2 wild-type RBD-XG005 and Omicron RBD-XG005. (A) Comparison the models of SARS-CoV-2 wild-type RBD-XG005 and Omicron RBD-XG005. Wild-type RBD and Omicron RBD were shown as ribbons and colored in light sky-blue and deep sky-blue, respectively. The XG005 IGHV and IGLV are colored in orange and yellow, respectively. Omicron mutation residues located in the XG005 epitope are shown as atom and colored in magenta. (B) The model of Omicron RBD-XG005. Omicron RBD is displayed in deep sky-blue. The XG005 epitope is colored in orange, and Omicron mutation residues within the interface located in XG005 epitope are shown as atoms and colored in magenta. (C and D) The detailed interfaces between SARS-CoV-2 wild-type RBD and XG005 and between Omicron RBD and XG005 (D). The red arrows emphasize the specific interactions between RBD and XG005. (E) Density maps of residues around the wild-type RBD-XG005 interface or Omicron RBD-XG005 interface. The Omicron mutations located in the XG005 epitope are labeled.

Based on the cryo-EM structure, the residues N450, V445, G447, N439 and Q506 of SARS-CoV-2 S protein were crucial for XG005 recognition, while SARS-CoV-2 VOCs bear no amino acid change on these residues. This is consistent with the overall high neutralizing potency of XG005 against all tested variants.

### Clonally-related neutralizing antibodies of XG005

It has been shown recently that a higher level of somatic hypermutation acquired in the months post-infection or by a vaccine booster shot provides some antibodies with greater neutralizing potency and breadth, suggesting that increased antibody diversity may improve antibody resistance to viral escape mutations (Gaebler et al., 2021; Muecksch et al., 2021; Sokal et al., 2021). However, XG005 was cloned from a donor early in convalescence and its somatic mutation level is low, with only 6 amino acid substitutions in both heavy and light chain V regions compared with germline sequences (Figure 5A). To understand the evolution process of XG005 for Omicron neutralization, we isolated three clonally-related antibodies of XG005 from the same donor (Zhou et al., 2021), and named them XG005a, XG005b and XG005c.

**Figure 5.**
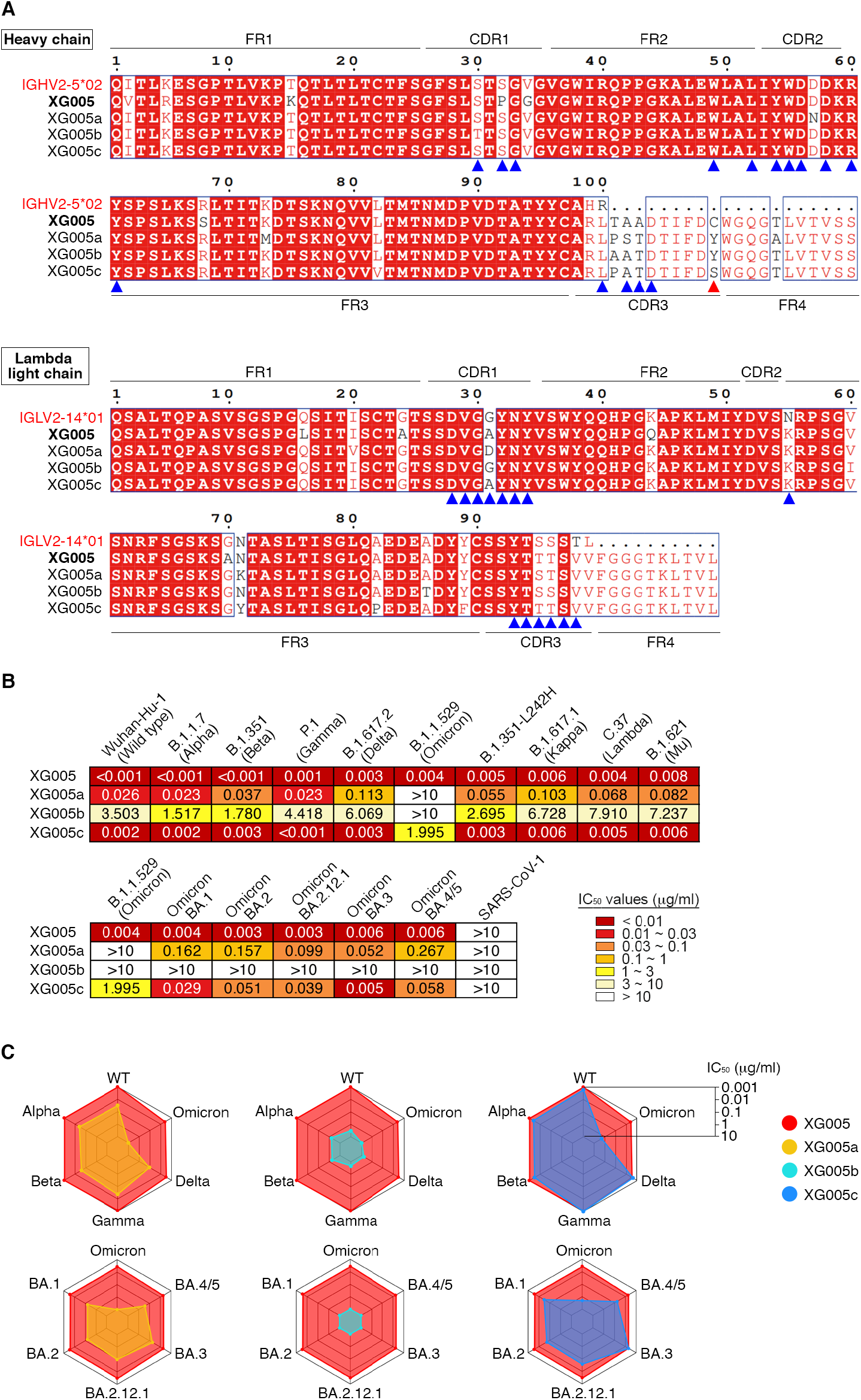
Clonally-related family members of XG005 exhibited striking disparity in neutralizing activity and breadth. (A) Amino acid sequence alignment for XG005 and its clonally-related family members, XG005a, XG005b and XG005c. IGHV2-5*02 and IGLV2-14*01 are the germline reference sequences assigned by IMGT/V-QUEST for IGHV and IGLV, respectively. Boxed red areas are shared among antibodies. The 15 amino acid residues in IGHV and 14 amino acid residues in IGLV involved in the recognition of the Omicron RBD are marked by blue arrowheads. The red arrowhead indicates the non-canonical cysteine C109 located in the CDR3 region of XG005 heavy chain. (B) Pseudovirus neutralization assays. IC_50_ values for four members measured against pseudoviruses of SARS-CoV-1, SARS-CoV-2 and its variants. Antibodies with IC_50_ values above 10 μg/ml were shown as >10 μg/ml. Mean of two independent experiments. (C) Spider charts for IC_50_ values of XG005 and its three family members.

XG005 and its three family members were all encoded by IGHV2-5/IGLV2-14 (Figure S5A). Sequence alignment between their heavy and light chains suggested a high similarity, including their CDR3 sequences of both heavy and light chains (Figure 5A). The levels of somatic hypermutation for all XG005 family members were low, and XG005a-c had even lower levels of somatic hypermutations compared with XG005 (Figure 5A).

We further evaluated their neutralization potency against a panel of pseudotyped viruses of SARS-CoV-2 variants. XG005 and its three family antibodies exhibited striking disparity in neutralizing activity and breadth (Figure 5B and S5B). Specifically, XG005b displayed minimal activity against most SARS-CoV-2 variant pseudoviruses, with IC_50_ values ranging from 1.517 to >10 μg/ml (Figure 5B and 5C). XG005a neutralized most variants, with IC_50_ values ranging from 0.023-0.267 μg/ml for all variants except Omicron, for which the IC_50_ was >10 μg/ml (Figure 5B and 5C). XG005c potently neutralized the majority of SARS-CoV-2 variant pseudoviruses (IC_50_ values of 0.001-0.058 μg/ml), but exhibited a partial loss of potency against Omicron (IC_50_ value of 1.995 μg/ml) (Figure 5B and 5C).

### Structural comparison for the key amino acid residues during antibody evolution

XG005 exhibited ultra-potent neutralizing activity against the B.1.1.529 (Omicron) pseudovirus, while none of the XG005 family members showed comparable activity. We further measured their ELISA binding activity against the S protein of B.1.1.529 (Omicron). As expected, similar binding activities were observed between XG005 and XG005c (ELISA AUC ∼30). XG005a had slightly reduced binding activity (ELISA AUC ∼26), while XG005b binding capacity was abolished by Omicron mutations (ELISA AUC ∼9) (Figure S5C).

To reveal the underlying molecular mechanism, we performed structural analysis and modeled the interactions between Omicron RBD and three XG005 family members (Figure 6A). The structural models of XG005a, XG005b, and XG005c were generated based on XG005 structure by SWISS-model (Waterhouse et al., 2018). The structures of all four XG005 family members were similar, with 15 amino acid residues in IGHV and 14 amino acid residues in IGLV involved in the recognition of the Omicron RBD (Figure 5A). Superimposed structural models showed that 8 of 11 key residues involved in the interaction were conserved among XG005 family members, including Y54, L52, R60, D56, and D58 in IGHV, and Y34, N33, and V98 in IGLV. However, although the residue D58 in IGHV was conserved among XG005 family members, this residue in XG005a, XG005b, and XG005c shifted away and damaged the hydrogen bond between N450 of Omicron RBD and D58 of antibody heavy chain, thus causing the reduced binding affinity against Omicron RBD (Figure 6B).

**Figure 6.**
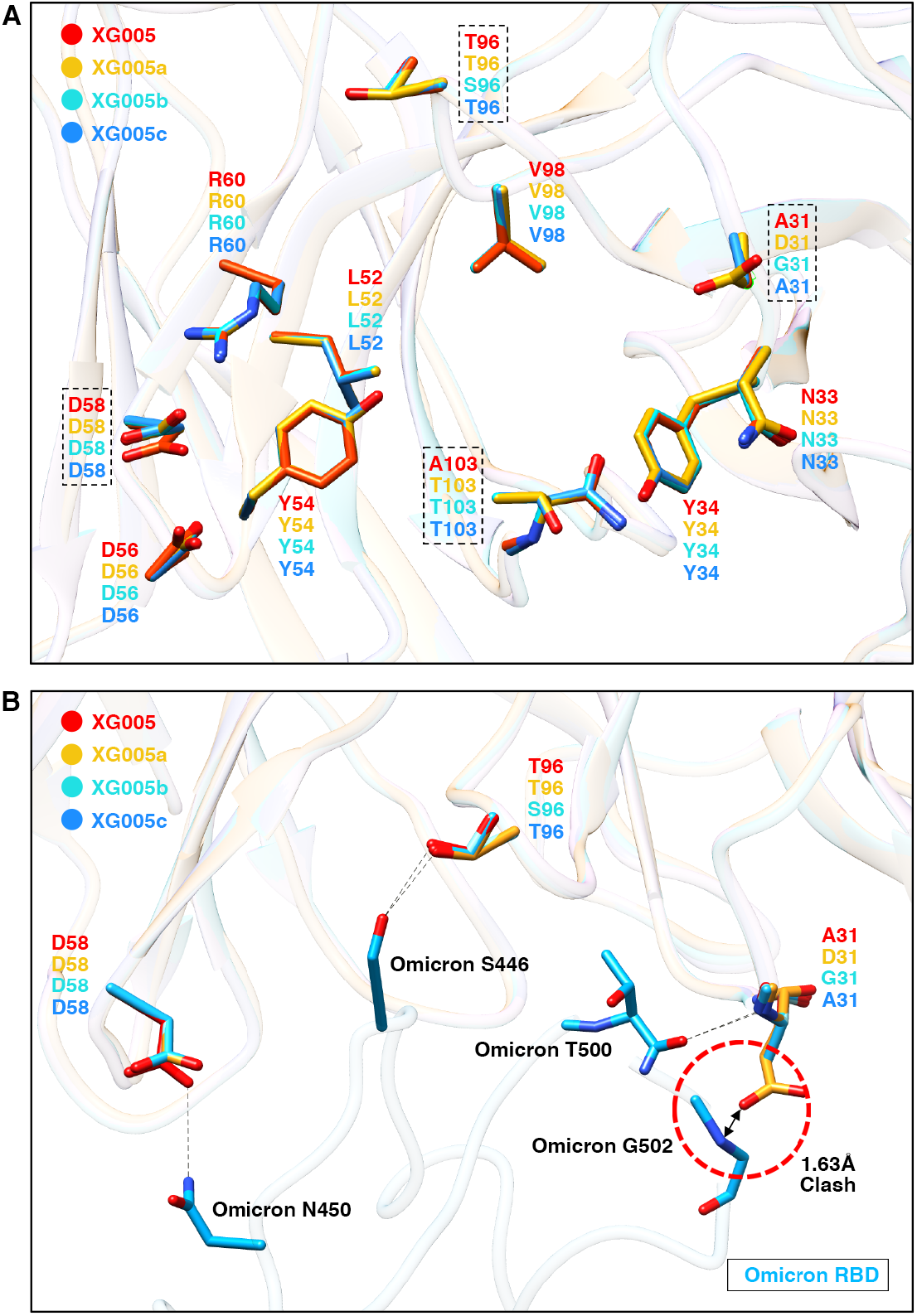
Structural comparison of XG005 family members revealed the key somatic mutations for broad and potent neutralization. (A) Structural comparison of XG005 (red ribbon), XG005a (yellow ribbon), XG005b (cyan ribbon) and XG005c (blue ribbon). The structural models of XG005a, XG005b and XG005c were generated by SWISS-MODEL. The residues involved in RBD binding are shown in sticks. The residues that might disturb the interactions between RBD and Fabs are emphasized by dashed squares. (B) The interfaces between Omicron RBD (deep sky-blue ribbon) and XG005/XG005a/XG005b/XG005c. The distinct key residues among XG005, XG005a, XG005b and XG005c were labeled. Hydrogen bonds are shown as black dashed lines. Red dashed circle highlights the clash between Omicron RBD and D31 of XG005a.

The other three key residues for Omicron RBD recognition were A103 in the heavy chain and T96/A31 in the lambda light chain of XG005 (Figure 6A and 6B). Both A103 in XG005 and T103 in other family members bound SARS-CoV-2 RBD with no difference. For the residue 96 of IGLV, the S96T mutation in XG005, XG005a and XG005c established a hydrogen bond between S446 of Omicron RBD and T96 of IGLV, while the lack of this somatic mutation in XG005b failed to do so (Figure 6B). In addition, the G31A mutation in XG005/XG005c IGLV was a key mutation for recognizing Omicron RBD. However, for XG005a, G31D mutation at this residue introduced a clash between Omicron RBD and D31 of XG005a, reducing XG005a’s binding affinity with Omicron RBD (Figure 6B). Together, these results provided a structural explanation that XG005 neutralized more potently than XG005c, and that XG005c neutralized better than XG005a and XG005b (Figure 5B and 5C).

### Engineered XG005 with reduced enhancement and extended half-life

XG005 was encoded by IGHV2-5/IGLV2-14. Similarly, as XG005, a well-known broad and potent neutralizing mAb, LY-CoV1404 (bebtelovimab), was also encoded by IGHV2-5/IGLV2-14 (Westendorf et al., 2022; Yuan et al., 2022). The cryo-EM structure of XG005 is extraordinarily comparable with that of LY-CoV1404 (Figure S6).

Our previous data showed that XG005 induced antibody-mediated viral entry and S protein-mediated membrane fusion through its interaction with Fc receptor (FcR), implying the potential side effect for antibody prophylaxis and therapy. (Liu et al., 2021; Zhou et al., 2021). As expected, LY-CoV1404 also induced in vitro antibody-dependent enhancement (ADE) of viral entry (Figure 7A). To eliminate its ADE of viral entry, we thus engineered XG005 Fc amino acids to reduce its FcR interactions (GRLR, G239R/L331R, or LALA, L237A/L238Amodifications). In vitro SARS-CoV-2 pseudovirus (ADE) assays (Zhou et al., 2021) showed that the engineered Fc variants of XG005 with GRLR or LALA substitutions induced no ADE effect in cultured Raji cells, while strong in vitro ADE effect was observed in Raji cells treated with wild-type XG005 (Figure 7B).

**Figure 7.**
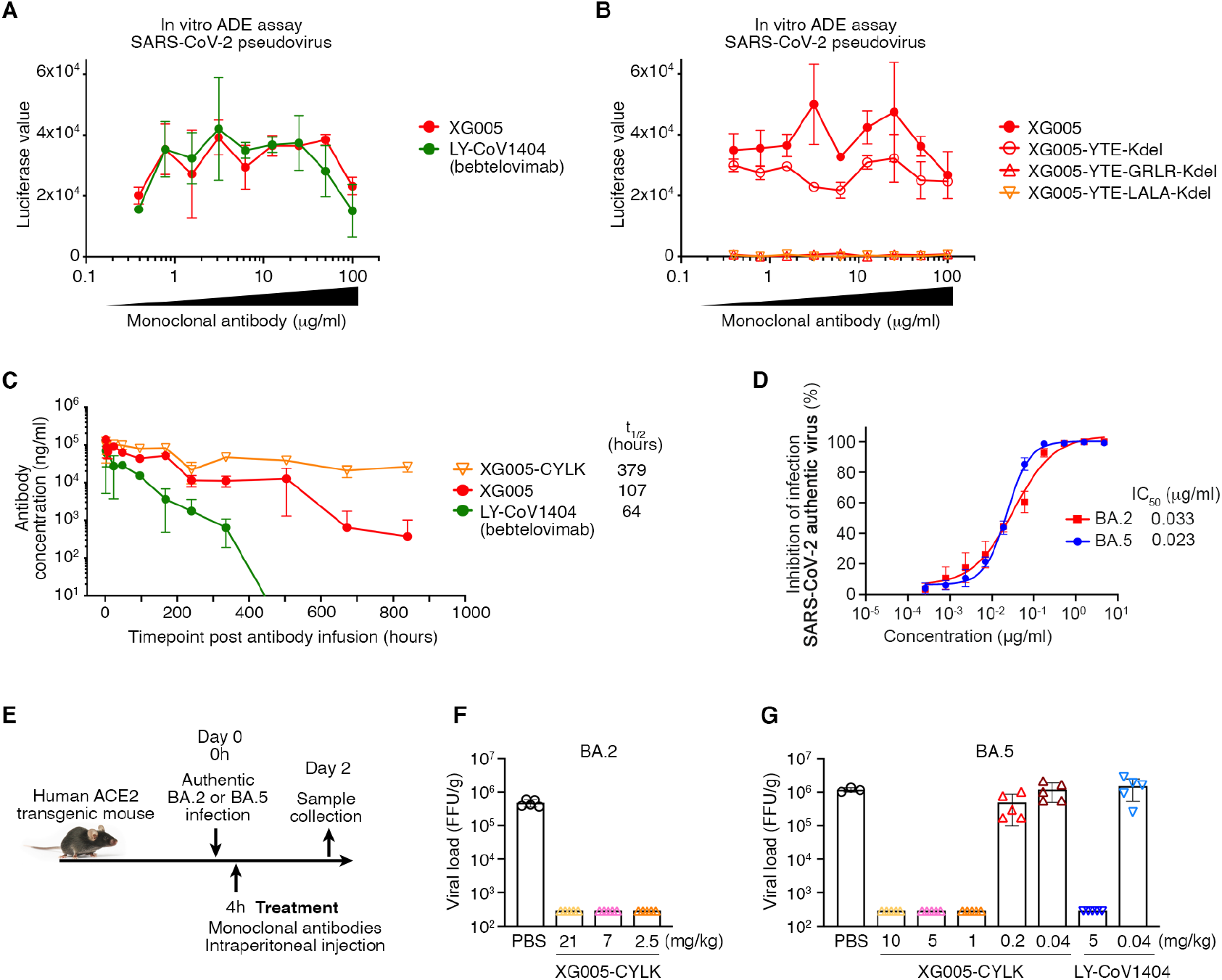
XG005-CYLK is therapeutic against BA.2 and BA.5 in vivo. (A) In vitro ADE effects induced by both XG005 and its counterpart LY-CoV1404 (bebtelovimab). In vitro ADE assays were performed in the Raji cells by using luciferase-expressing SARS-CoV-2 pseudovirus. The presence of various dilutions of antibodies induced distinct levels of luciferase signal, while the luciferase signal without adding any antibody was almost zero. (B) No ADE effect induced by the GRLR and LALA version, but not YTE version, of the Fc-engineered XG005 antibodies. (C) Pharmacokinetics of single-dose mAbs, XG005, XG005-C109S-YTE-LALA-Kdel (XG005-CYLK), and LY-CoV1404, in transgenic mice, C57BL/6JSmoc, which expressed human neonatal Fc receptor (hFcRn). (D) XG005 potently neutralizes authentic SARS-CoV-2 BA.2 and BA.5 viruses. The in vitro neutralization assays were repeated at least twice. (E) iagram of antibody treatment protocols for human ACE2 transgenic mice intranasally challenged with BA.2 or BA.5 viruses. (F-G) Virus titers in lung tissues of mice collected at two days after BA.2 (F) or BA.5 (G) viral infection. Data are presented as mean ± SD. Each group contains three to five individual mice.

XG005 had a non-canonical cysteine (C109, red arrowhead in Figure 5A) in the CDR3 region of XG005 heavy chain. To avoid the potential aggregation and antibody instability triggered by this non-canonical cysteine through intramolecular scrambling or intermolecular disulfide formation (Buchanan et al., 2013), we substituted the cysteine at position 109 with tyrosine (Y) or serine (S) residues, the corresponding amino acid residues in XG005b and XG005c, respectively (Figure 5A). Both substitutions (XG005-C109Y and XG005-C109S) showed no reduction in neutralizing activity against SARS-CoV-2 variants (Figure S7).

We further engineered XG005 Fc domain to extend the antibody half-life (YTE, M255Y/S257T/T259E, or LS, M431L/N437S) and to reduce antibody heterogeneity (Kdel, deletion of the C-terminal lysine in Ig heavy chain). Pharmacokinetic analyses in humanized FcRn transgenic mouse showed that XG005 had a longer half-life than LY-CoV1404, and that the YTE substitution in XG005 significantly extended its serum half-life (Figure 7C). Moreover, we performed in vitro neutralization assays to ensure that none of these Fc substitutions affect the in vitro neutralizing activity of XG005 (Figure S8).

Together, to facilitate and improve therapeutic use, we engineered XG005, reduced its ADE effect, increased its half-life, optimized the antibody production and quality, and finally developed XG005-C109S-YTE-LALA-Kdel (XG005-CYLK) for the following therapeutic evaluation in vivo.

### Therapeutic activity of XG005-CYLK

We first confirmed the neutralizing activity of XG005-CYLK using authentic SARS-CoV-2 viruses BA.2 and BA.5 (Figure 7D). As shown, significant inhibitory activities against BA.2 and BA.5 infection were observed for XG005-CYLK, with IC_50_ values of 0.033 μg/ml and 0.023 μg/ml, respectively.

We next sought to evaluate the therapeutic activity of XG005-CYLK in an established human ACE2 transgenic mouse model. Mice were intranasally challenged with BA.2 or BA.5 virus using 1×10^5^ focus forming units (FFU), and, four hours post infection, intraperitoneally treated with a single dose of antibody XG005-CYLK or the same volume of PBS as control (Figure 7E). Two days post inoculation, the viral loads in the lungs reached 3.7∼6.1×10^5^ FFU for BA.2 and 1.03∼1.37×10^6^ FFU for BA.5 in the control groups of mice treated with PBS (Figure 7F and 7G). Compared with the control groups, a single dose of XG005-CYLK (2.5 mg/kg, 7 mg/kg or 21 mg/kg for BA.2; 1 mg/kg, 5 mg/kg or 10 mg/kg for BA.5) efficiently reduced the viral loads by more than 1,000-fold in the lungs (Figure 7F and 7G). However, decreased levels of XG005-CYLK (0.2 mg/kg and 0.04 mg/kg for BA.5) were not sufficient to suppress the lung viral loads (Figure 7G).

Collectively, these results suggest that rare B cells elicited by just wild-type SARS-CoV-2 infection retained broad neutralization against the currently circulating SARS-CoV-2 variants, and the corresponding mAbs could be engineered as potent therapeutics.

## DISCUSSION

In this article, we examined the binding capacity and neutralizing activity of 45 monoclonal antibodies (mAbs) isolated from a convalescent individual who donated blood in April 2020. Among them, the XG005 monoclonal antibody showed potent and broad neutralizing activity against all variants, including the BA.2, BA.2.12.1, and BA.4/5 subvariants. Treatment experiments in mice with engineered XG005 alone showed efficient viral-controlling effect in vivo. Therefore, the high threshold against virus escape provided by an antibody cocktail would also be achieved by a single mAb alone.

Cryo-EM structure explained how XG005 avoided immune escape and maintained neutralization potency. Although many bNAbs against Omicron bound to an outer surface of RBD (Fang et al., 2022; Nutalai et al., 2022; Pinto et al., 2020), XG005 recognized the receptor-binding motif (RBM), which bound to ACE2 receptor and is highly mutated in the Omicron subvariants (VanBlargan et al., 2022). However, distinct from the immune escape of most RBM-targeting antibodies, XG005 delicately avoided significant loss of neutralization despite the three Omicron amino acid mutations (N440K, G446S and N501Y) located in the XG005-recognizing epitope. Newly formed hydrogen bonds and salt bridges simultaneously rescued the loss of hydrogen bonds between XG005 and Omicron S protein (Figure 4). This effective compensation mechanism plays an important role for recognizing various SARS-CoV-2 variants.

XG005 was an IGHV2-5/IGLV2-14-encoded RBD antibody, while antibodies LY-CoV1404 (bebtelovimab) (Westendorf et al., 2022), 2-7 (Kramer et al., 2021) and XGv265 (Wang et al., 2022a) were also IGHV2-5/IGLV2-14-encoded (Yuan et al., 2022). All of these mAbs retained their neutralizing activity against SARS-CoV-2 variants, especially against Omicron and its sub-lineages. Comparison of the cryo-EM structures of both XG005 and LY-CoV1404 revealed very high level of similarity, including the RBD interfaces and the key amino acid residues for RBD interaction (Figure S6). Compared with IGHV1-58/IGKV3-20-encoded RBD-binding public antibody clonotype, the IGHV2-5/IGLV2-14-encoded RBD antibodies with increased breadth of neutralization, are perhaps the most cross-neutralizing public antibody clonotype (Yuan et al., 2022).

In our study, three other family members of XG005 (XG005a, XG005b, and XG005c) we cloned from the same expanded B cell clone of the same donor, showed significant reduction of neutralization potency and breadth. Nevertheless, sequence comparison showed only very little difference of these mAbs compared with XG005. Structural remodeling suggested the key amino acid residues on XG005 during antibody evolution for its neutralizing activity against Omicron subvariants. Considering that there was still no Omicron variant during blood donation in April 2020, these results suggest that XG005 was the rare product of random somatic hypermutation in germinal centers. Similarly, LY-CoV1404 was also cloned from a convalescent individual in early 2020 (Westendorf et al., 2022). Interestingly, the third dose of vaccine booster shot with wild-type SARS-CoV-2 facilitated the generation of potent bNAbs against VOCs and Omicron subvariants (Wang et al., 2022a). Together, we conclude that the exposure to wild-type SARS-CoV-2 or its surface protein is sufficient to elicit bNAbs against recently emerged or even future SARS-CoV-2 variants.

## MATERIALS AND METHODS

### ELISA

To perform ELISA, 96-well microplates were coated with antigen proteins (10 μg/ml) in phosphate-buffered saline (PBS) (50 μl per well) overnight at 4℃. Antigen proteins were S-ECD protein of SARS-CoV-2 Wuhan-Hu-1 (wild-type) and its related variants, including Alpha (B.1.1.7), Beta (B.1.351), Gamma (P.1), Delta (B.1.617.2) and Omicron (B.1.1.529). These coated plates were then blocked with PBS containing 2% bovine serum albumin (BSA) (200 μl per well). After blocking, the plate was incubated with the first antibody (eight dilutions with a maximum concentration of 10 μg/ml, 3-fold serially diluted) in PBS (50 μl per well) for one hour at room temperature. After wash, the second antibody (goat anti-human IgG conjugated with HRP, Thermo Fisher Scientific) in PBS (50 μl per well) was added to each well for another one-hour incubation, and then detection was performed. To evaluate the antigen-binding capacity, we calculated the area under the curve (AUC) for each purified recombinant IgG1 monoclonal antibody using PRISM software as previously reported (Wang et al., 2020).

#### Generation of SARS-CoV-1/2 pseudoviruses

Pseudotyped viruses of SARS-CoV-1, SARS-CoV-2 and SARS-CoV-2-related variants were generated as described previously (Xia et al., 2020; Zhou et al., 2021). We first constructed the S-protein expression plasmids pcDNA3.1-SARS-CoV-2-S or pcDNA3.1-SARS-CoV-1-S. The S protein amino acid sequences for SARS-CoV-2 wild-type and variants were provided (Table S2). We then co-transfected the constructed pcDNA3.1 plasmids with the backbone plasmid of pNL4-3.Luc.R-E into HEK293T cells. Two days later, we collected the cell supernatants containing pseudoviruses and stored them at -80℃ for in vitro neutralization assays.

#### In vitro pseudotyped virus-based neutralization assay for SARS-CoV-2

We performed the in vitro pseudovirus neutralization assays as previously described (Xia et al., 2020; Zhou et al., 2021). We first examined the generated pseudotyped viruses by infecting Huh-7 cells and measuring luciferase activity to determine the virus concentration. We then aliquoted the concentrated virus stock and stored it at -80℃. To perform the in vitro neutralization experiments, we seeded Huh-7 cells in 96-well plates (10^4^ cells per well) and serially diluted (1:3) the overexpressed monoclonal antibodies (maximum concentration 10 μg/ml) for nine dilutions in total. We mixed and incubated the antibody soup and concentrated pseudovirus soup for 30 minutes at 37℃, and then added them into the Huh-7 cells for twenty-four hours of incubation. We then replated the cell supernatant with fresh DMEM containing 10% FBS and collected cells after 36 hours of cell culture. Finally, we lysed the cultured cells and measured luciferase activity using a Firefly Luciferase Assay Kit (Promega, USA) and a microplate reader (Infinite M200PRO, Switzerland) following the manufacturer’s instructions. Due to the dramatic variation of the absolute luciferase values, we calculated the relative luminescence values by normalizing them to pseudovirus-only control wells. The IC_50_ values by neutralization assays were calculated by nonlinear regression analysis in PRISM software.

#### In vitro neutralization assay using authentic BA.2 and BA.5 viruses

Experiments including viral amplification and viral infection were conducted in a Biosafety Level 3 (BSL-3) laboratory. The authentic BA.2 and BA.5 viruses were amplified and titered in Vero-E6 cells using the plaque assay. The in vitro neutralization assay was performed as described previously (Liu et al., 2021; Zhou et al., 2021). Different concentrations of mAbs were mixed with the authentic BA.2 or BA.5 viruses for 1 hour before adding onto cultured cells. Twenty-four hours later, the cells were fixed and subjected to immunostaining assay to determine the cell infection rate.

#### Antibody cloning and production

Single B cell-based antibody amplification and sequence analysis were performed as previously reported (Wang et al., 2020; Zhou et al., 2020). Briefly, we performed the reverse transcription and nested PCR amplification for the sorted single B cells (Zhou et al., 2021). We analyzed all the Sanger sequencing results of heavy and light chains and identified the V(D)J gene and CDR3 sequences using IMGT/V-QUEST (Brochet et al., 2008) and/or IgBLAST (Ye et al., 2013). For antibody expression, we transiently transfected HEK293F cells with heavy/light chain plasmids and harvested supernatants seven days later for antibody purification.

#### Expression and purification of SARS-CoV-2 Omicron S trimer

The vector of Omicron S ectodomain with HexaPro mutations, “GSAS” substitution at furin cleavage site (residues 682-285) and a C-terminal T4 fibritin trimerization was constructed as previously reported (Li et al., 2022) and transfected into HEK293F cells for expression.

After 72 hours, the supernatants were harvested and filtered for affinity purification by Histrap HP (GE Healthcare). The protein was then loaded onto a Superose 6 increase 10/300 column (GE Healthcare) in 20 mM Tris pH 8.0, 200 mM NaCl.

#### Cryo-EM sample preparation

Purified SARS-CoV-2 Omicron S at 0.60 mg/mL was mixed with XG005 antibody by a molar ratio of 1:1.7 and incubated for 10 minutes on ice. A 3 μl aliquot of the sample was loaded onto a freshly glow-discharged holey amorphous nickel-titanium alloy film supported by 400 mesh gold grids. The sample was frozen immediately in liquid ethane using Vitrobot IV (FEI/Thermo Fisher), with 2 s blot time and -3 blot force and 10 s wait time.

#### Cryo-EM data collection and image processing

Cryo-EM data were collected on a Titan Krios microscope (Thermo Fisher) operated at 300 kV. Movies were captured with a K3 summit direct detector (Gatan) after a GIF quantum energy filter (Gatan) setting to a slit width of 20 eV. Automated data acquisition was carried out with SerialEM software (Mastronarde, 2005) through beam-image shift method (Wu et al., 2019).

Movies were taken in the super-resolution mode at a nominal magnification 81,000×, corresponding to a physical pixel size of 1.064 Å, and a defocus range from −1.2 μm to −2.5 μm. Each movie stack was dose-fractionated to 40 frames with a total exposure dose of about 58 e^−^/Å^2^ and exposure time of 3s.

A total of 6,503 movie stacks was motion corrected using MotionCor2 (Zheng et al., 2017) within RELION (Zivanov et al., 2018). Parameters of contrast transfer function (CTF) were estimated by using Gctf (Zhang, 2016). All micrographs then were manually selected for further particle picking upon astigmatism, defocus range and estimated resolution.

Remaining 6,098 good images were imported into cryoSPARC (Punjani et al., 2017) for further patched CTF-estimating, blob-picking and 2D classification. Several good 2D classes were used as templates for template-picking. After 2D classification of particles from template-picking was finished, all good particles from blob-picking and template-picking were merged and deduplicated, subsequently being exported back to RELION through pyem package (Asarnow, 2019).

Total 2,028,032 particles were extracted at a box-size of 320 and rescaled to 160, then carried on one round of 3D classification in RELION. Only good classes were selected, yielding 1,594,120 particles. These particles were performed other rounds of 3D classification to get different states of trimer. Finally, three main states with clear Fabs were selected out, and their corresponding particles were separately re-extracted (unbinned, 1.064 Å/pixel) and auto-refined, then CTF-refined and polished. 153,541 of state 1 (1-RBD-up with 2 Fabs) was yielding a map at 3.62 Å, 124,608 of state 2 (1-RBD-up with 3 Fabs) was yielding a map at 3.74 Å, and 616,627 of state 3 (2-RBD-up with 3 Fabs) was yielding a map at 3.24 Å.

To get more clear structural information of interface between RBD with Fab, we carried on local 3D-classification focused on the best pair of RBD and Fab from state 3. In final, 313,560 particles were subtracted and exported to cryoSPARC to do local refinement, yielding a 2.99 Å local map.

The reported resolutions above are based on the gold-standard Fourier shell correlation (FSC) 0.143 criterion. The above procedures of data processing are summarized (Figure S3 and S4). These sharpened maps were generated by DeepEMhancer (Sanchez-Garcia et al., 2021) and then “vop zflip” to get the correct handedness in UCSF Chimera (Pettersen et al., 2004) for subsequent model building and analysis.

#### Model building and refinement

For model building of SARS-CoV-2 Omicron S XG005 complex, the SARS-CoV-2 Omicron S trimer model and the antibody model generated by swiss-model (Waterhouse et al., 2018) were fitted into the map using UCSF Chimera and then manually adjusted with COOT (Emsley et al., 2010). The interface between RBD and Fab region was refined against the local refinement map and then docked back into the into global refinement trimer maps. Several iterative rounds of real-space refinement were further carried out in PHENIX (Afonine et al., 2018). Model validation was performed using MolProbity. Figures were prepared using UCSF Chimera and UCSF ChimeraX (Pettersen EF, 2021).

The cryo-EM maps and the coordinates of SARS-CoV-2 Omicron S complexed with XG005 have been deposited to the Electron Microscopy Data Bank (EMDB) and Protein Data Bank (PDB) with accession numbers EMD-33744 and PDB 7YD0 (state 1, UDD with two Fabs), EMD-33742 and PDB 7YCY (state 2, UDD with three Fabs), EMD-33743 and PDB 7YCZ (state 3, UDU with three Fabs), and EMD-33745 and PDB 7YD1 (Local refinement).

#### Human ACE2 transgenic mice and in vivo studies

Mouse experiments were conducted in a Biosafety Level 3 (BSL-3) laboratory in Guangzhou Custom technology center. Transgenic mice with human ACE2 overexpression (hACE2-Tg) were randomly assigned to distinct groups. A single administration of mAbs (or an equal volume of PBS as a negative control) was administered intraperitoneally 4 hours after all mice were intranasally challenged with 10^5^ PFU BA.2 or BA.5 authentic viruses. Mouse body weight was monitored, and lungs were collected two days post-infection to determine the live viral loads in lungs by the focus forming assay (FFA).

#### Pharmacokinetic analysis

Transgenic mice (C57BL/6JSmoc) with human neonatal Fc receptor (hFcRn) overexpression (Vendor: Shanghai Model Organisms Center, China) were used to evaluate the pharmacokinetic profiles of mAbs XG005, XG005-CYLK, and LY-CoV1404 (bebtelovimab). Twenty-seven mice were randomly assigned into three groups and a single dose of mAbs (10 mg/kg) were administrated based on their body weights. The serums samples were collected on different time points, including -1-day pre-infusion and 2, 4, 8, 24, 48, 96, 168, 240, 336, 504, 672, 840 hours post-infusion. Sample analysis was conducted utilizing validated ELISA methods. Sample concentration data was collected on the INFINITE 200 PRO plate reader and processed using INFINITE 200 PRO Software (2013) Tecan. Pharmacokinetic parameters were evaluated using a non-compartmental approach with Phoenix WinNonlin software (Version 8.0.0.3176, Pharsight, CA).

## ACKNOWLEDGMENTS

We thank Center of Cryo-Electron Microscopy at Fudan University for the support on cryo-EM data collection, and Guangzhou Custom Technology Center for the support of in vivo challenge study in a Biosafety Level 3 (BSL-3) laboratory. We thank Bo Chen, Xulong Feng, Xinyi An, Miaomiao Wang, Yongpeng Xu, Qingyu Yang at Advaccine Biopharmaceuticals Suzhou Co. Ltd. for the help of cell-based and pharmacokinetics assays. We also thank Dr. Xiangxi Wang at Institute of Biophysics, Chinese Academy of Sciences for providing the S proteins of several SARS-CoV-2 variants for ELISA assays. This work was supported by National Key Research and Development Program (2021YFA1301400), National Natural Science Foundation of China (31872730 and 32070947), Ministry of Science and Technology of China (2021YFC2302500). Project was also supported by Shanghai Municipal Science and Technology Major Project (ZD2021CY001) and by Guangzhou Laboratory (SRPG22-003). This work was supported by funding from the Intramural Research Program, National Institutes of Health, National Cancer Institute, Center for Cancer Research. The content of this publication does not necessarily reflect the views or policies of the Department of Health and Human Services, nor does mention of trade names, commercial products, or organizations imply endorsement by the U.S. Government. The content is solely the responsibility of the authors and does not necessarily represent the views of any of the funding agencies or individuals.

## AUTHOR CONTRIBUTIONS

Conceptualization, Q.W.; Investigation, J.W., Z.C., Y.G., Z.W., J.W., B.Y.C., Y.Z., Y.H., W.Z., M.X., W.J., X.Z., A.H., A.X., J.H., and S.X.; Software, J.W., W.Z., and Z.C.; Formal Analysis, J.W., W.Z., Y.Z., B.W., Z.W., J.W., B.Y.C., L.Z., L.S., and Q.W.; Writing – Original Draft, Q.W.; Writing – Review & Editing, J.W., W.Z, C.T.M, B.Y.C., B.W., L.S., and Q.W.; Visualization, J.W., W.Z., L.S., and Q.W.; Supervision, B.W., F.W., L.Z., L.S. and Q.W.; Funding Acquisition, B.W., F.W., L.Z., L.S., and Q.W.

## COMPETING INTERESTS

A patent application encompassing aspects of this work has been filed with Q.W. and L.Z. listed as an inventor. Other authors have no conflicts of interest to declare.

## SUPPLEMENTARY FIGURES AND LEGENDS

**Figure S1.**
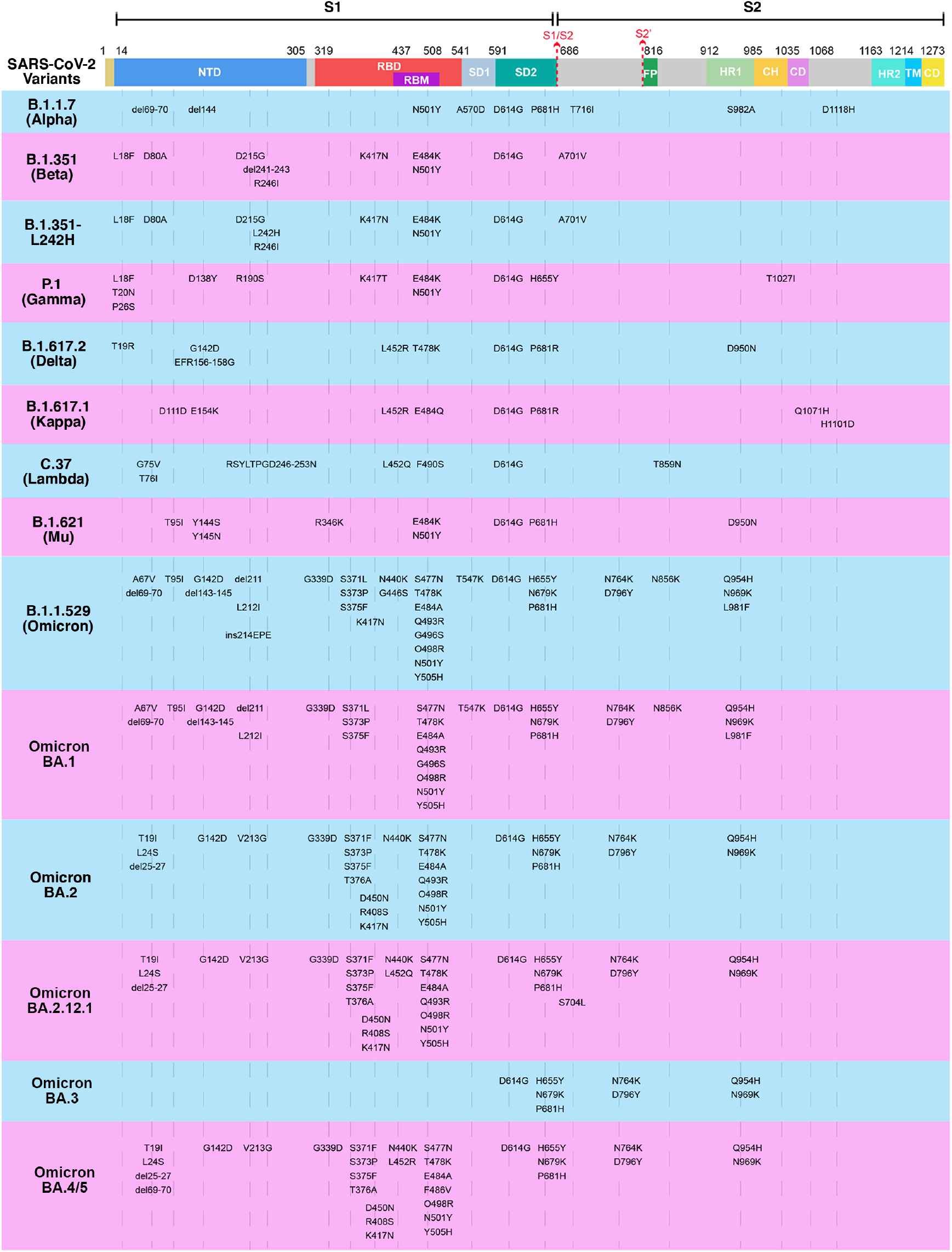
S protein mutations within SARS-CoV-2 variants. Key spike mutations found in the SARS-CoV-2 variants, compared with Wuhan-Hu-1 (wild-type), are denoted. These variants include five SARS-CoV-2 VOCs: B.1.1.7 (Alpha), B.1.351 (Beta), P.1 (Gamma), B.1.617.2 (Delta) and B.1.1.529 (Omicron); five Omicron variants: BA.1, BA.2, BA.2.12.1, BA.3 and BA.4/5; and four other variants: B.1.351-L242H, B.1.617.1 (Kappa), C.37 (Lambda) and B.1.621 (Mu).

**Figure S2.**
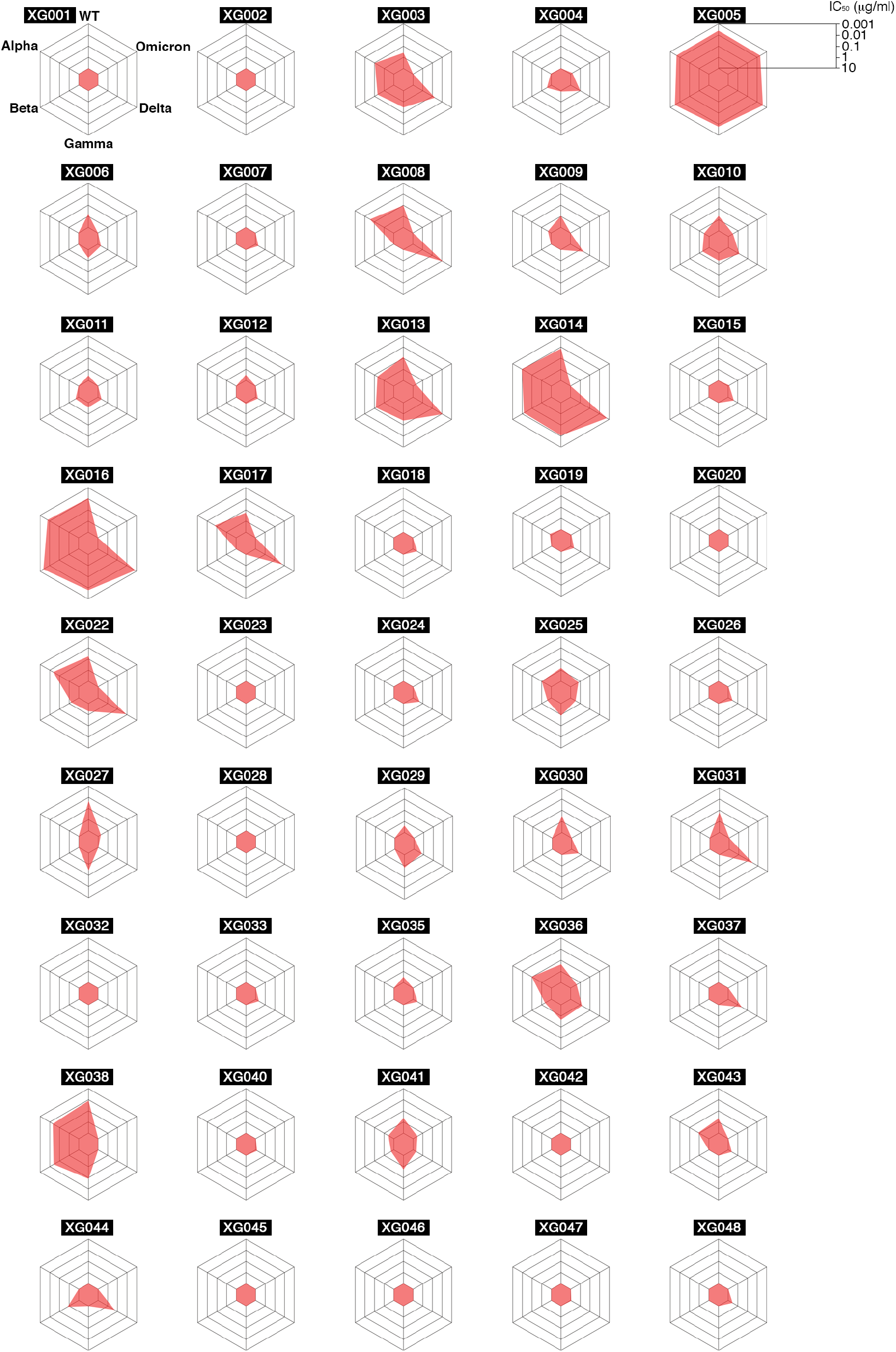
Spider charts for IC_50_ values of 45 monoclonal antibodies. IC_50_ values against Wuhan-Hu-1 (wild-type), B.1.1.7 (Alpha), B.1.351 (Beta), P.1 (Gamma), B.1.617.2 (Delta) and B.1.1.529 (Omicron) pseudoviruses were measured for all 45 tested antibodies isolated from a convalescent donor (Zhou et al., 2021). Antibodies with IC_50_ values (mean of two independent experiments.) above 10 μg/ml were shown as 10 μg/ml.

**Figure S3.**
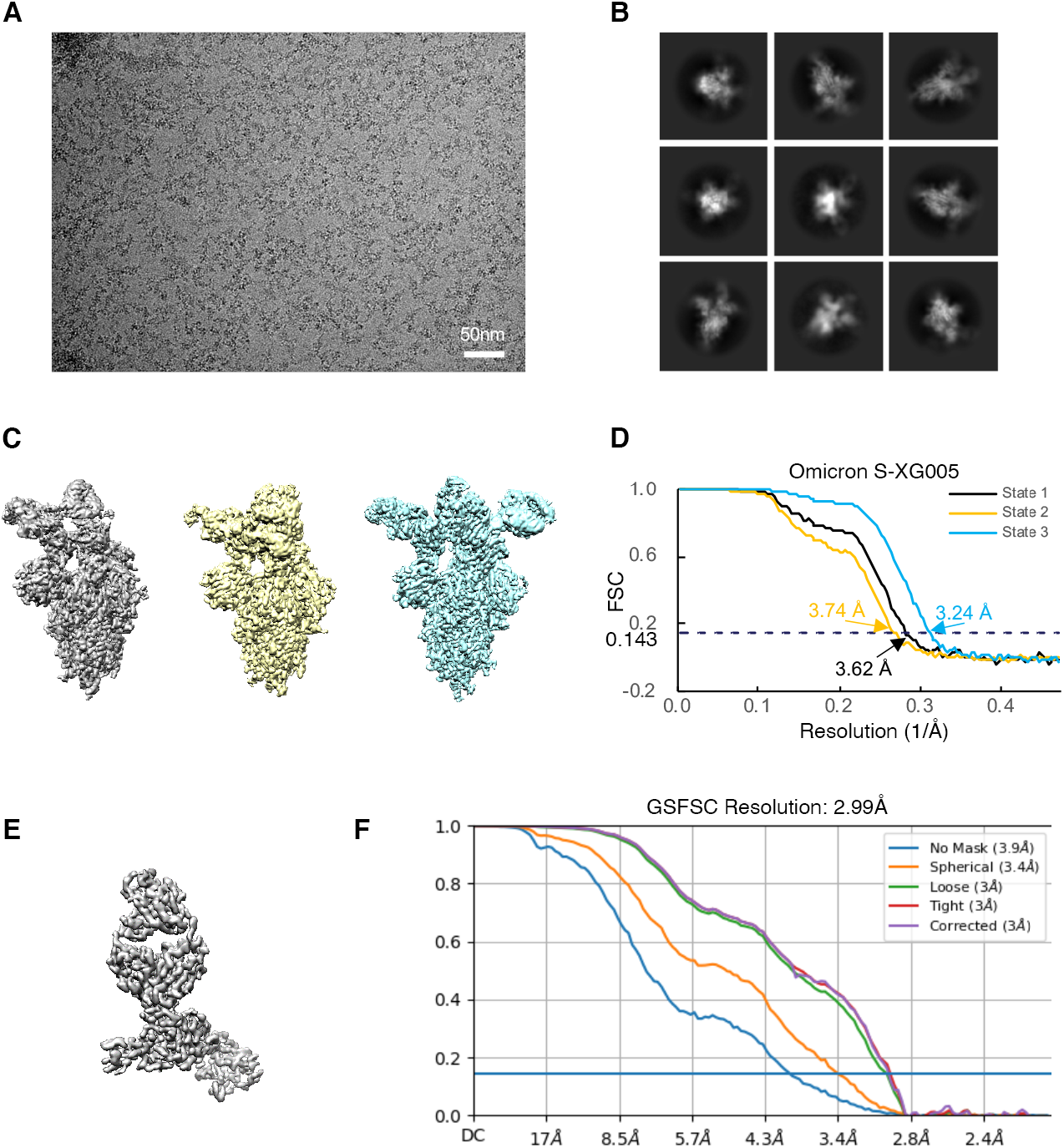
Cryo-EM data collection and processing of Omicron S-XG005 complex (OS-XG005). (A) Representative electron micrograph. (B) 2D classification results of OS-XG005. (C) The reconstruction maps of the complex structures at three states. (D) Gold-standard Fourier shell correlation curves (FSC) for each structure. The 0.143 cut-off is indicated by a horizontal dashed line. (E) The local refined map of the interface between RBD and Fab region. (F) FSC of local refinement of RBD-XG005 Fab region obtained from cryoSPARC.

**Figure S4.**
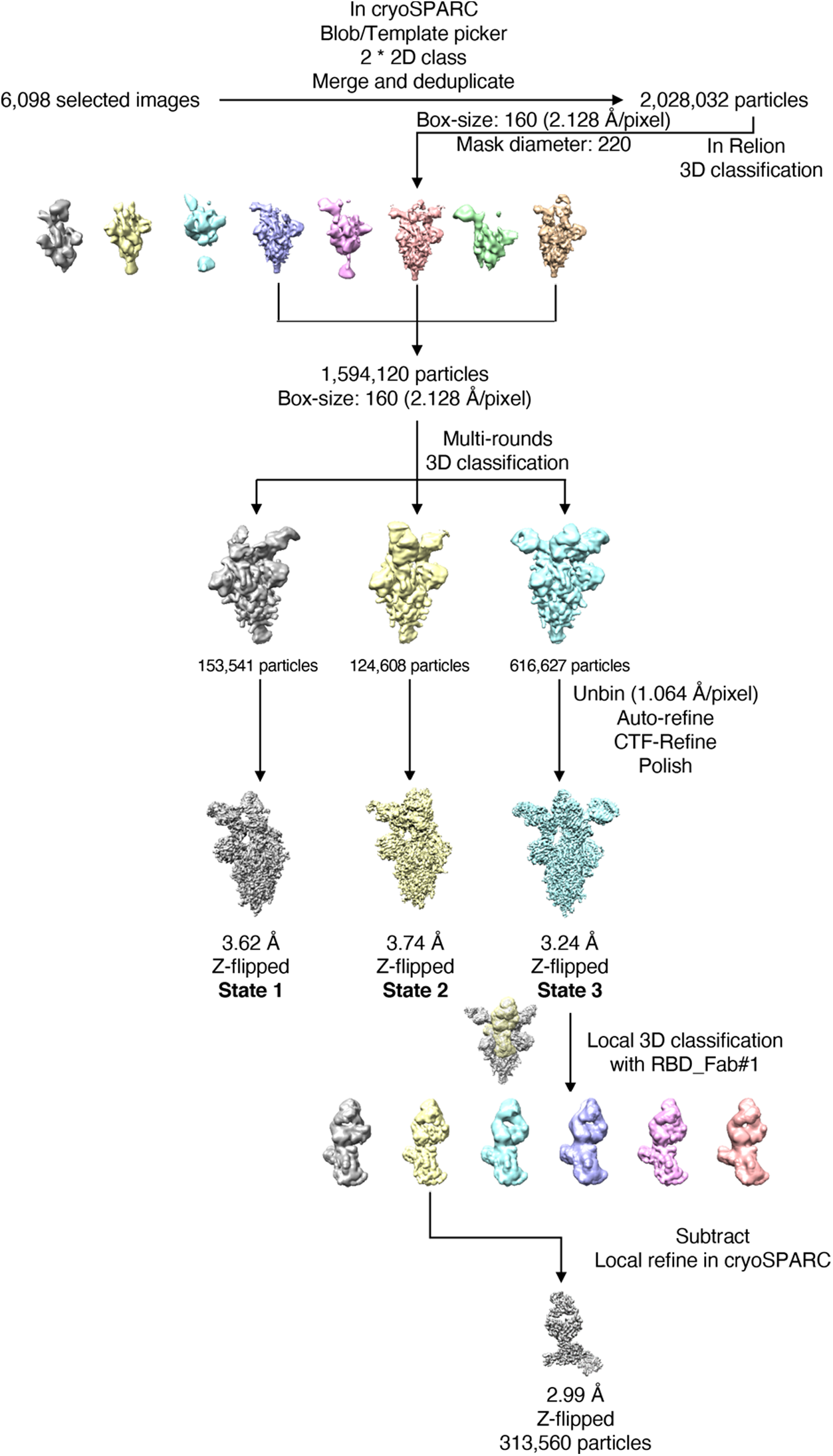
Data processing flowchart of OS-XG005 complex.

**Figure S5.**
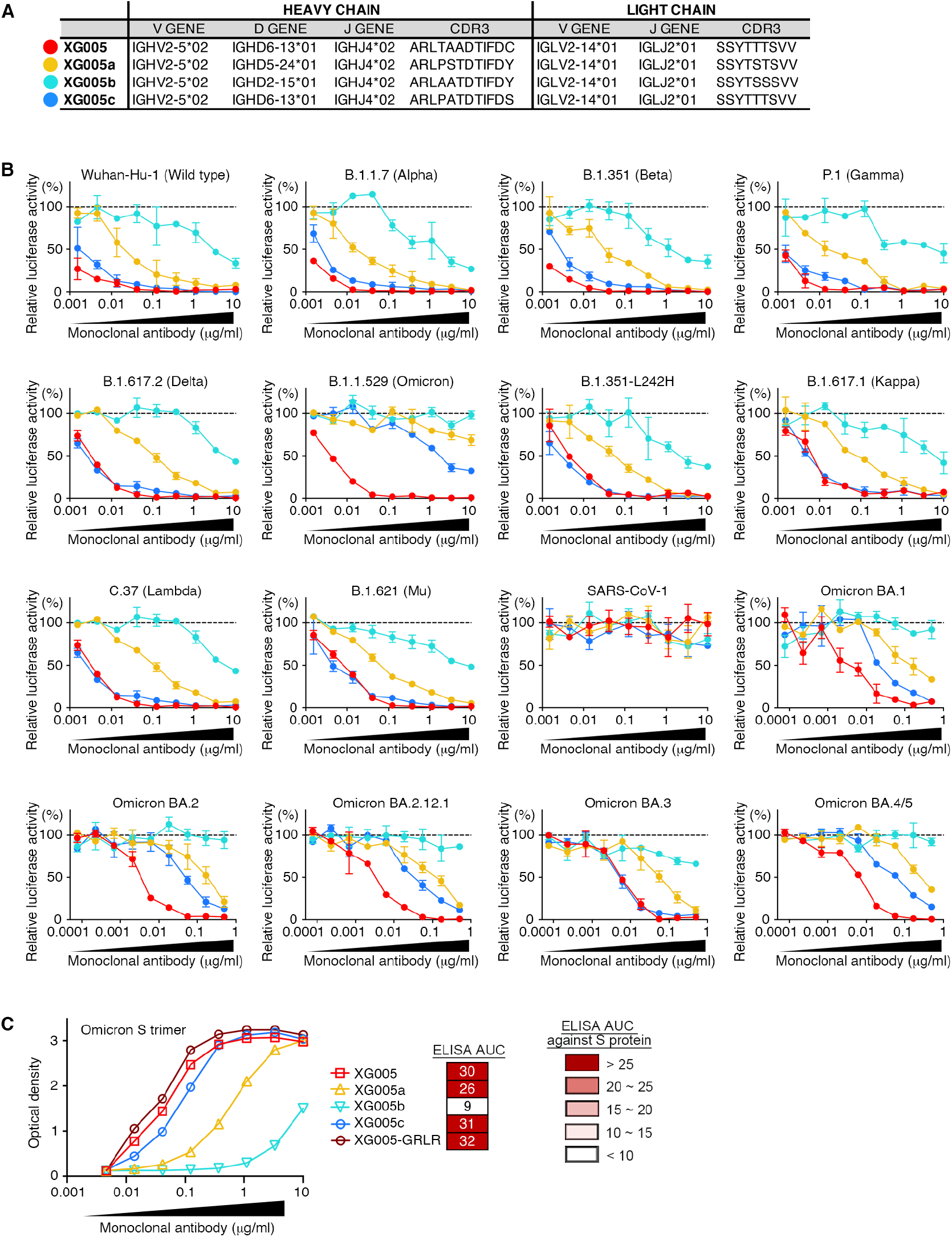
In vitro pseudovirus neutralization assays for XG005 family members. (A) V(D)J assignments for the XG005 clone. IMGT/V-QUEST was used to assign the V, D, J genes and CDR3 sequences for their Ig heavy and light chains. (B) Neutralization potency of XG005 family members. Luciferase-based pseudoviruses were used for in vitro infection. Dashed line represents no antibody control. All experiments were repeated at least twice, presented as mean ± SEM. (C) Dramatically distinct binding capacity against Omicron S protein by XG005 family members. ELISA assays to determine the antibody binding capacity against Omicron S proteins. ELISA area under the curve (AUC) values were calculated. XG005c showed similar level of binding activity of XG005, while those of XG005b and XG005c dramatically reduced.

**Figure S6.**
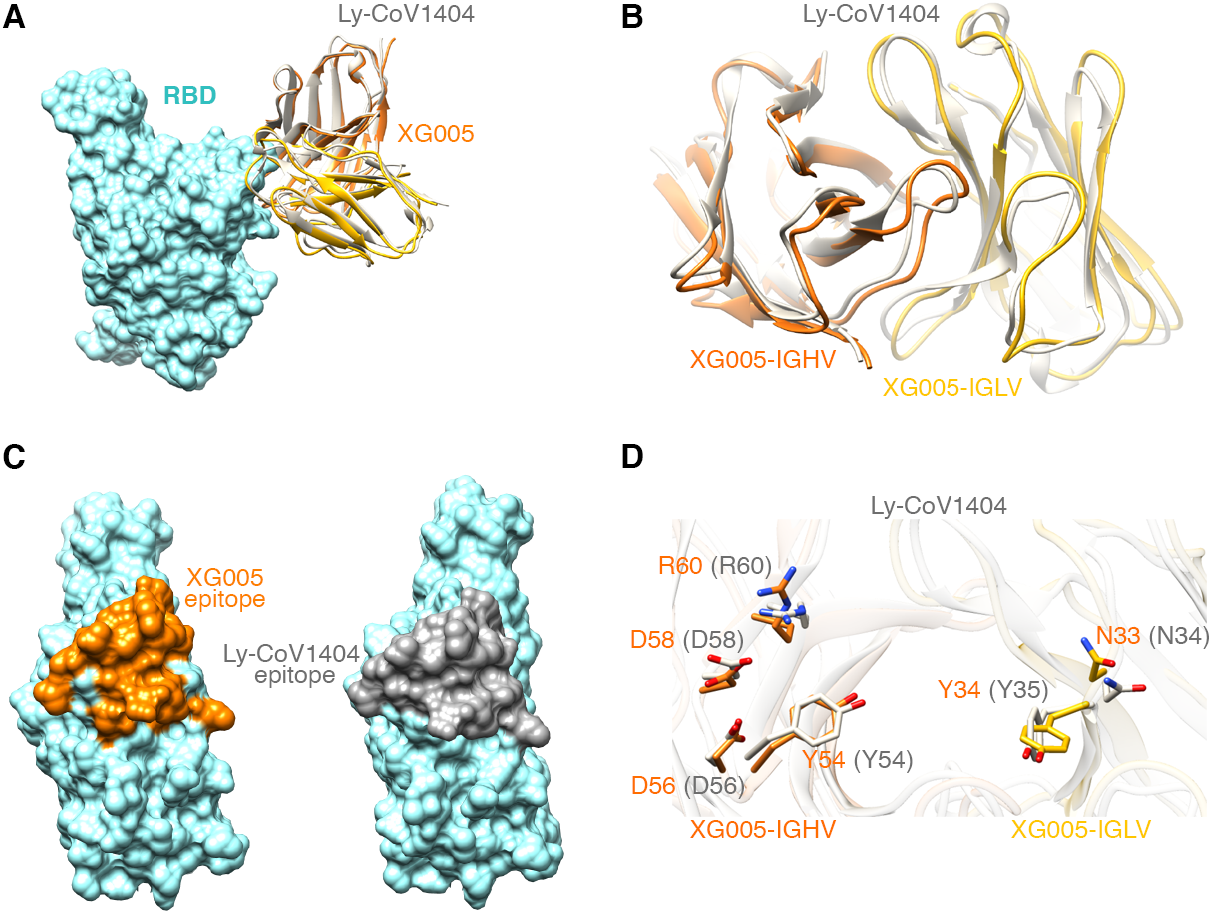
Structural comparison between XG005 and LY-CoV1404. (A) Comparison the models of wild-type RBD complexed with XG005 and LY-CoV1404. RBD is displayed in sky-blue surface; XG005 heavy and light chains are shown in orange and yellow ribbons, respectively, while LY-CoV1404 is shown in gray. (B) A close view of XG005 and LY-CoV1404. (C) Surface representation of RBD showing the interfaces of XG005 (orange) and LY-CoV1404 (dark gray), respectively. (D) Comparison of the key residues of XG005 and LY-CoV1404 involved in the RBD interaction. Residues of XG005 and LY-CoV1404 are labeled in orange and gray, respectively.

**Figure S7.**
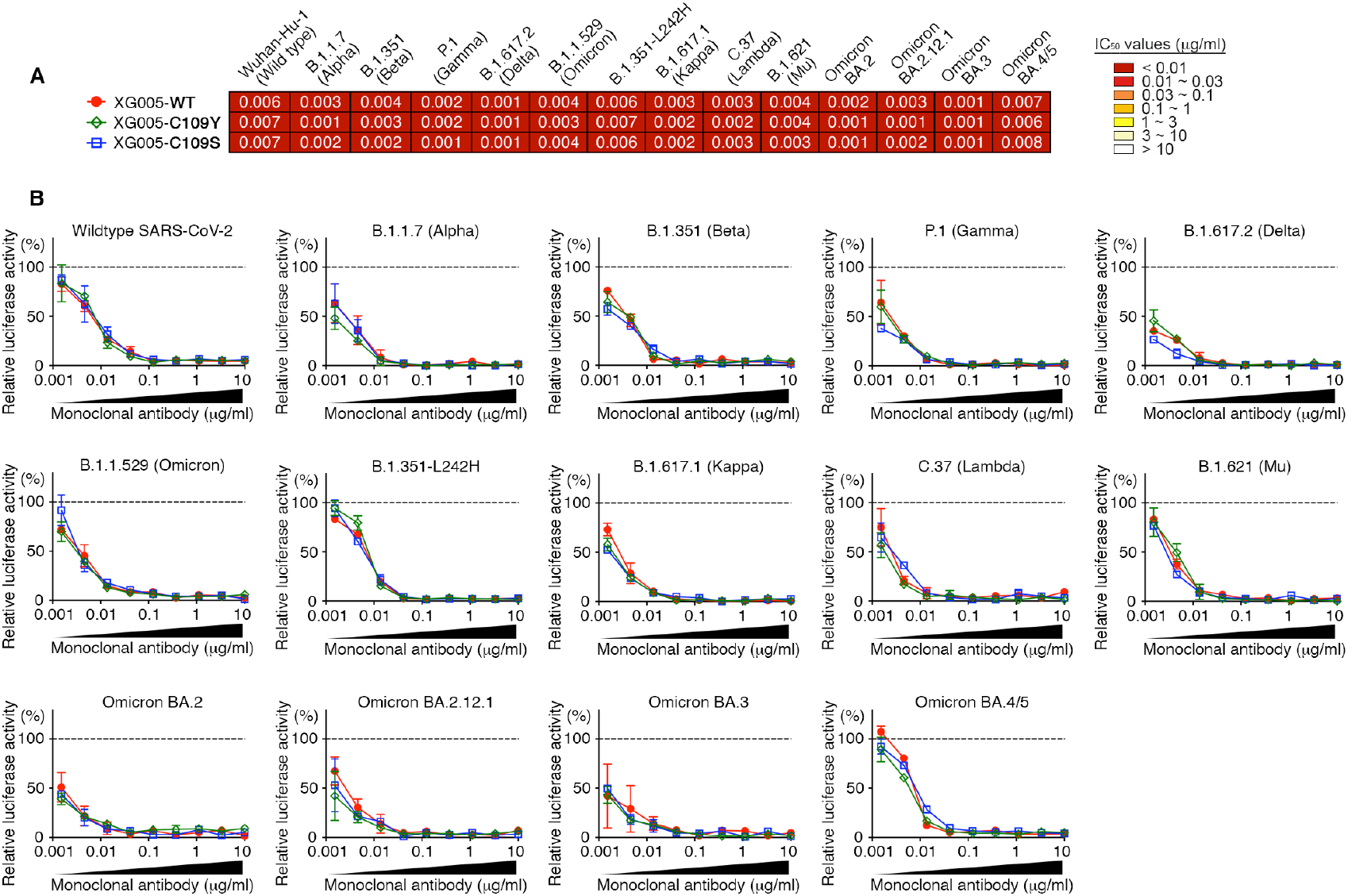
Engineered C109 variants of XG005 maintain neutralization potency. (A) IC_50_ values for XG005-WT, XG005-C109Y, and XG005-C109S measured against pseudoviruses of SARS-CoV-1, SARS-CoV-2 and its variants. (B) Pseudovirus neutralization assays using different concentrations of XG005-WT, XG005-C109Y, and XG005-C109S. Mean of two independent experiments.

**Figure S8.**
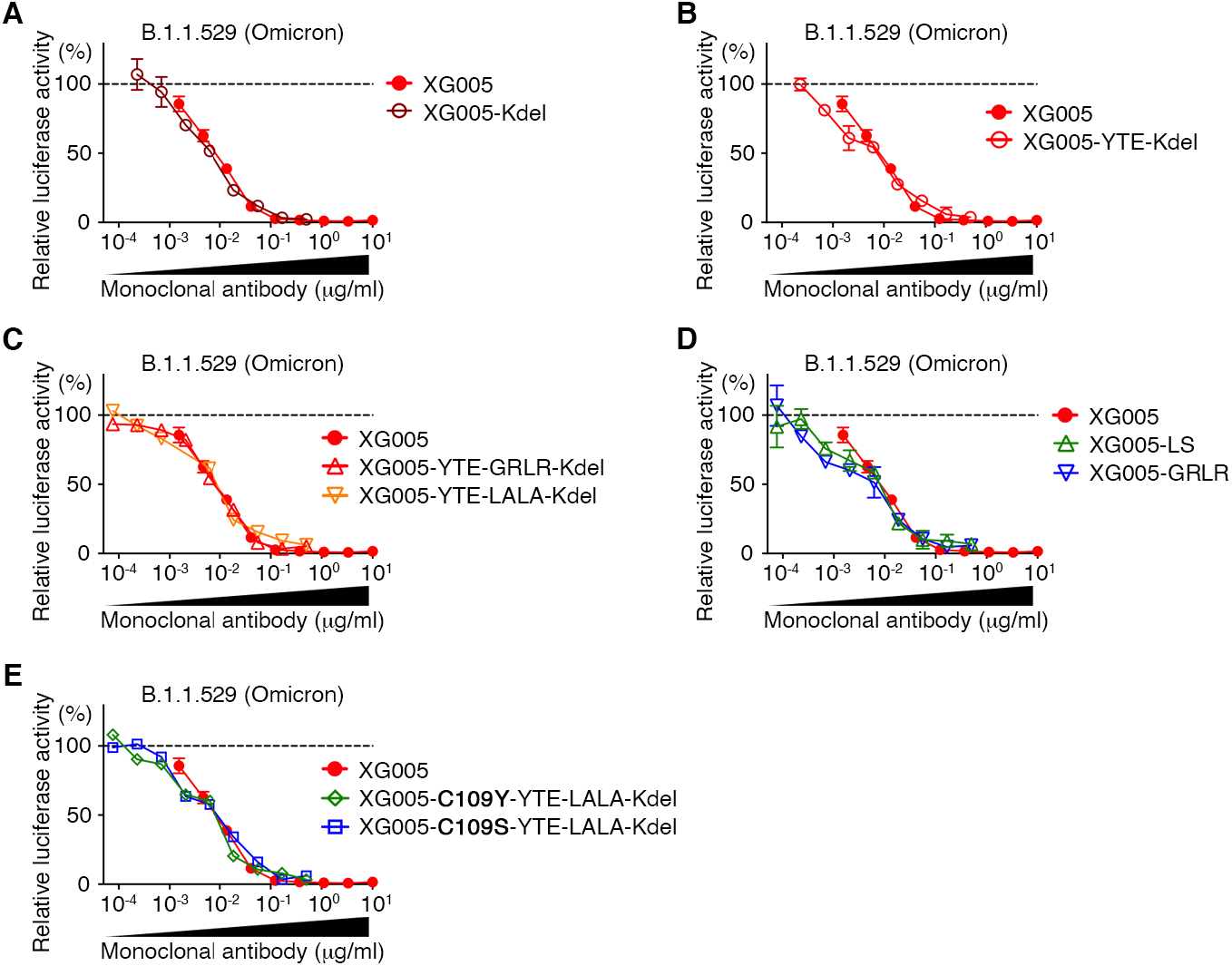
Engineered Fc variants of XG005 maintain neutralization potency. (A-D) Various engineered Fc variants of XG005 maintain the in vitro neutralizing activities against B.1.1.529 (Omicron) pseudoviruses. Kdel: mAb mutant with the deletion of heavy chain C-terminal lysine (A). YTE: mAb mutant with triple mutations M255Y, S257T and T259E in the Fc domain (B and C). LS: mAb mutant with M431L and N437S mutations in the Fc domain (D). Both YTE and LS substitutions result in an increase in its binding to human FcRn and a prolonged serum half-life of the antibody. GRLR: mAb mutant with G239R and L331R mutations in the Fc domain (C and D). LALA: mAb mutant with L237A and L238A mutations in the Fc domain (C). Both GRLR and LALA substitutions abrogate the antibody binding to FcγRs and eliminate the ADE effect.

**Table S1.** Cryo-EM data collection and refinement statistics of the Omicron S-XG005 complex.

|  | State 1<br>(UDD with<br>two Fabs) | State 2<br>(UDD with<br>three Fabs) | State 3<br>(UDU with<br>three Fabs) | RBD+Fab<br>XG005 |
| --- | --- | --- | --- | --- |
| **Data collection and processing** |  |  |  |  |
| Magnification | 81,000 |  |  |  |
| Voltage (kV) | 300 |  |  |  |
| Total dose (e-/Å<sup>2</sup>) | 58 |  |  |  |
| Defocus range (µm) | -1.2 to -2.5 |  |  |  |
| Pixel size (Å) | 1.064 |  |  |  |
| Symmetry imposed | C1 |  |  |  |
| Final particles (no.) | 153,541 | 124,608 | 616,627 | 313,560 |
| Map resolution (Å) | 3.62 | 3.74 | 3.24 | 2.99 |
| **Refinement** |  |  |  |  |
| **R.m.s. deviations** |  |  |  |  |
| Bond lengths (Å) | 0.002 | 0.002 | 0.002 | 0.002 |
| Bond angles (°) | 0.558 | 0.509 | 0.517 | 0.479 |
| **Validation** |  |  |  |  |
| MolProbity score | 2.40 | 2.24 | 2.23 | 2.46 |
| Clashscore | 7.16 | 5.99 | 6.11 | 6.30 |
| Rotamer outlier (%) | 5.83 | 4.53 | 4.85 | 5.46 |
| **Ramachandran plot** |  |  |  |  |
| Favored (%) | 93.64 | 93.72 | 94.54 | 90.05 |
| Allowed (%) | 6.36 | 6.28 | 5.44 | 9.95 |
| Disallowed (%) | 0.00 | 0.00 | 0.00 | 0.00 |
| **EMDB** | EMD-33744 | EMD-33742 | EMD-33743 | EMD-33745 |
| **PDB** | 7YD0 | 7YCY | 7YCZ | 7YD1 |

**Table S2.**
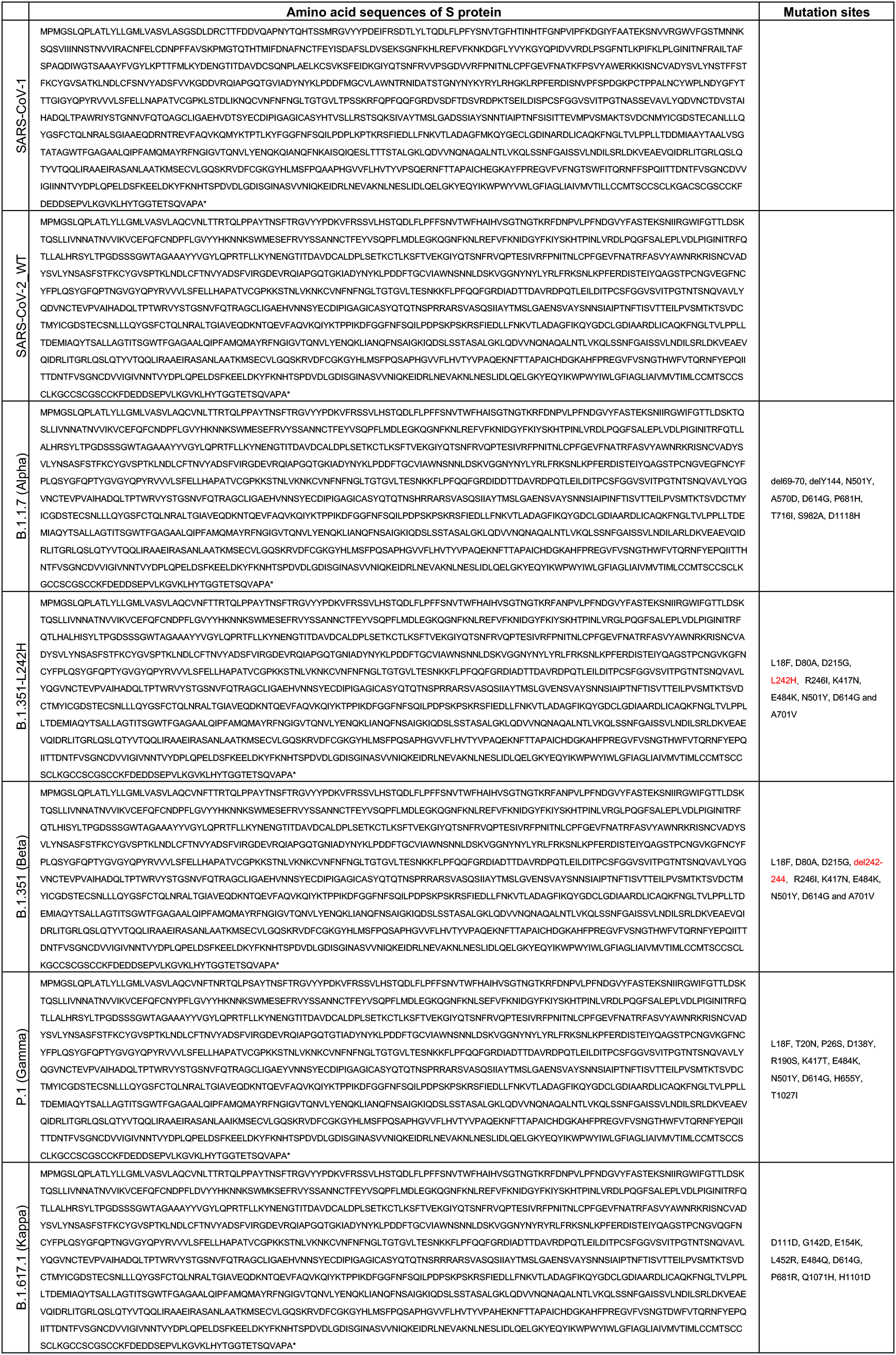
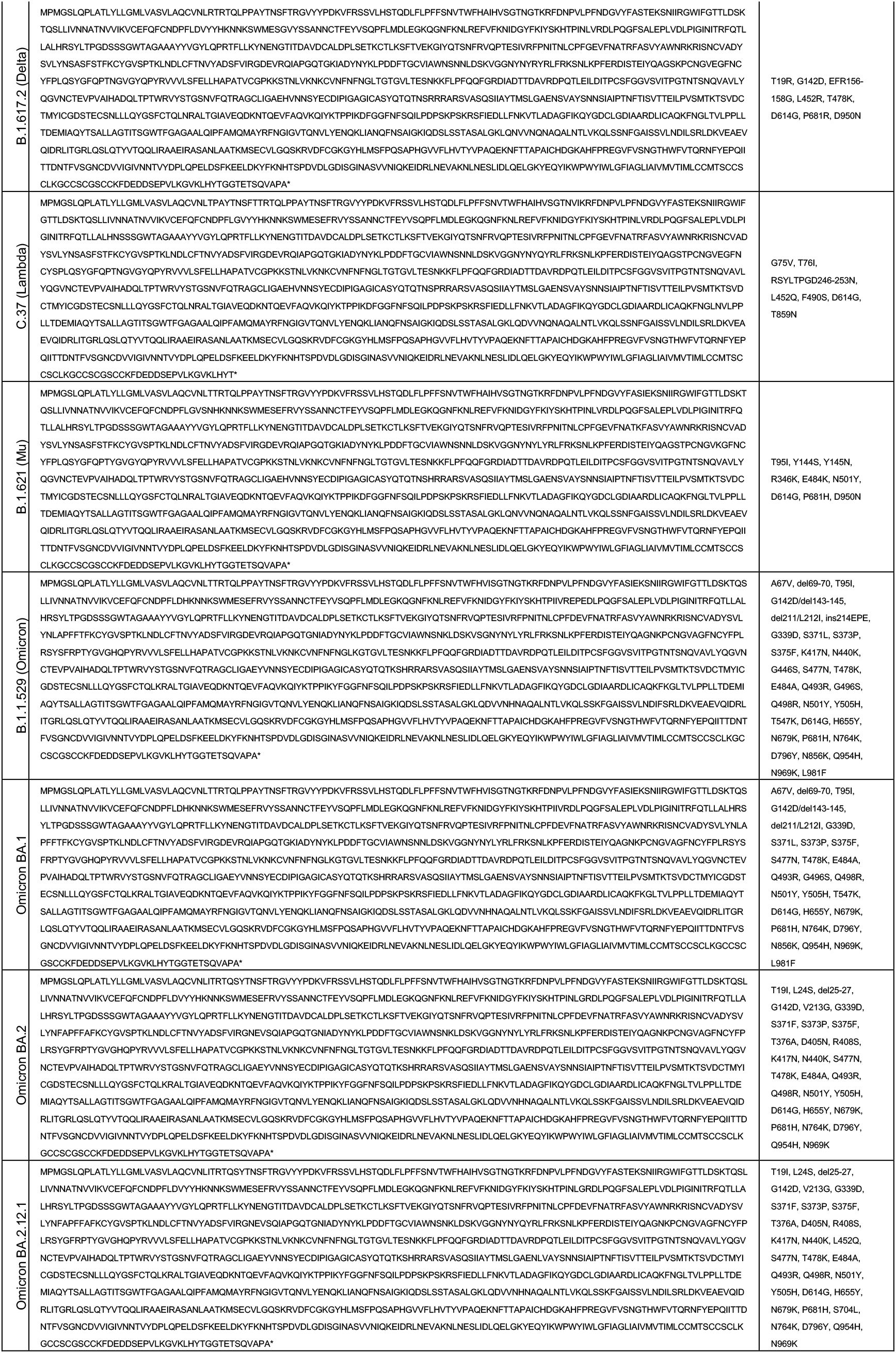
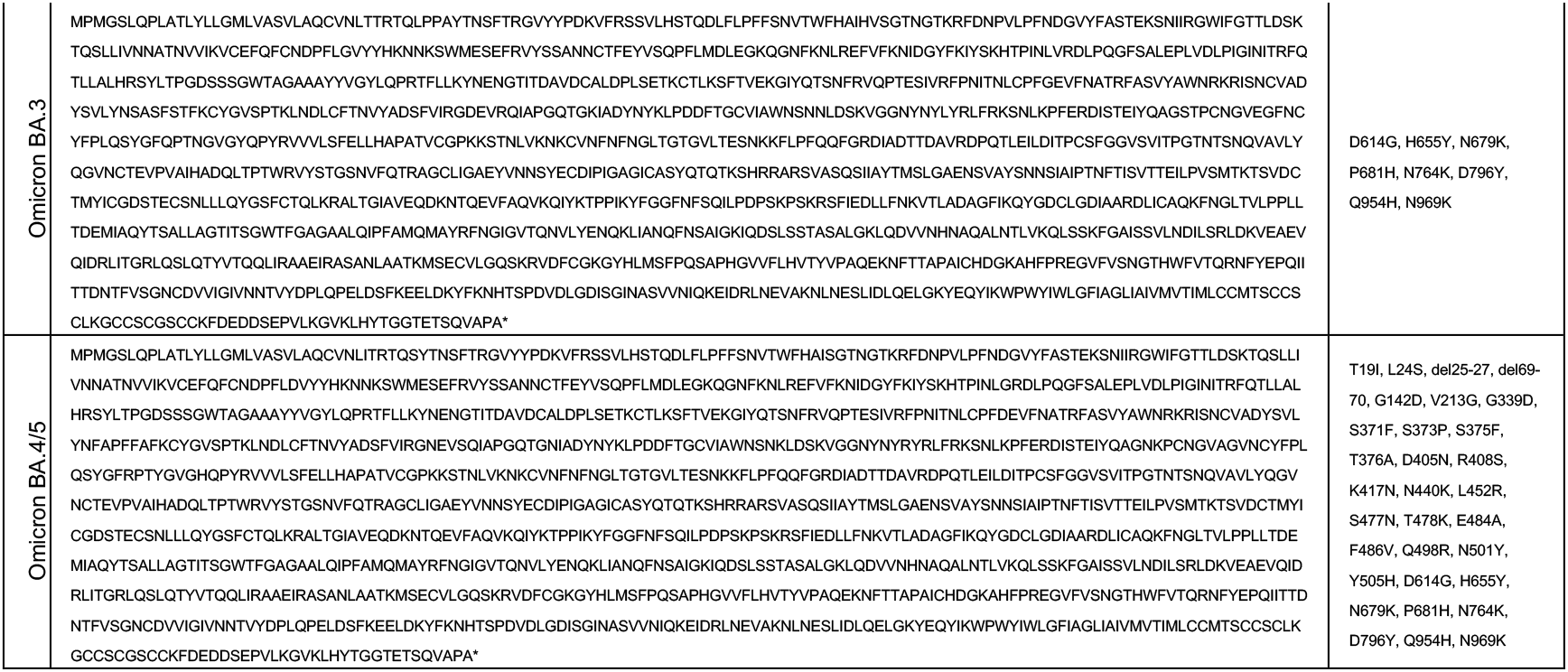
Amino acid sequences of the S protein of various pseudotyped viruses used for in vitro neutralization assays.

